# High Shear Stress Reduces ERG Causing Endothelial-Mesenchymal Transition and Pulmonary Arterial Hypertension

**DOI:** 10.1101/2024.02.02.578526

**Authors:** Tsutomu Shinohara, Jan-Renier Moonen, Yoon Hong Chun, Yannick C. Lee-Yow, Kenichi Okamura, Jason M. Szafron, Jordan Kaplan, Aiqin Cao, Lingli Wang, Shalina Taylor, Sarasa Isobe, Melody Dong, Weiguang Yang, Katherine Guo, Benjamin D Franco, Cholawat Pacharinsak, Laura J. Pisani, Shinji Saitoh, Yoshihide Mitani, Alison L. Marsden, Jesse M. Engreitz, Jakob Körbelin, Marlene Rabinovitch

## Abstract

Pathological high shear stress (HSS, 100 dyn/cm^2^) is generated in distal pulmonary arteries (PA) (100-500 μm) in congenital heart defects and in progressive PA hypertension (PAH) with inward remodeling and luminal narrowing. Human PA endothelial cells (PAEC) were subjected to HSS versus physiologic laminar shear stress (LSS, 15 dyn/cm^2^). Endothelial-mesenchymal transition (EndMT), a feature of PAH not previously attributed to HSS, was observed. H3K27ac peaks containing motifs for an ETS-family transcription factor (ERG) were reduced, as was ERG-Krüppel-like factors (KLF)2/4 interaction and ERG expression. Reducing ERG by siRNA in PAEC during LSS caused EndMT; transfection of ERG in PAEC under HSS prevented EndMT. An aorto-caval shunt was preformed in mice to induce HSS and progressive PAH. Elevated PA pressure, EndMT and vascular remodeling were reduced by an adeno-associated vector that selectively replenished ERG in PAEC. Agents maintaining ERG in PAEC should overcome the adverse effect of HSS on progressive PAH.

## INTRODUCTION

Computational modeling revealed levels of abnormally elevated high shear stress (HSS, 100 dyn/cm^2^) in the peripheral pulmonary arteries (PAs) of a congenital heart defect (CHD) with increased pulmonary blood flow and associated pulmonary arterial hypertension (APAH) such as a ventricular septal defect^1^. Similar levels of pathological HSS have been computed in the distal pulmonary circulation of patients with idiopathic (I) pulmonary arterial hypertension (PAH) as the vessels undergo inward remodeling resulting in elevated pulmonary vascular resistance (PVR) to flow^2, 3^. Pathologic HSS has been studied as a factor that induces PAH in animal models, most recently in a rat with an aorto-caval arteriovenous (AV) shunt injected with the toxin monocrotaline, in which there is severe progressively severe disease^4^. Mice with an AV shunt have also been created and they develop increasing PAH attributed to vascular remodeling over a period of eight weeks^5, 6^ as do postnatal lambs in which an aorto-pulmonary shunt is created in fetal life^7^.

However, the mechanism whereby HSS initiates and induces progression of pulmonary vascular changes, that ultimately cause irreversible elevation in PVR and severe PAH, is not known. Relative to static conditions, physiologic laminar shear stress (LSS, 15 dyn/cm^2^) is characterized by an increase in Krüppel-like factor 2 and 4 (KLF2/4), in their interaction with SWItch/Sucrose Non-Fermentable (SWI/SNF) chromatin remodelers, and in induction of protective genes such as endothelial nitric oxide synthase *(NOS3),* bone morphogenetic protein receptor 2 *(BMPR2),* mothers against decapentaplegic 5 *(SMAD5)* and repression of genes associated with vasoconstriction and smooth muscle cell (SMC) proliferation such as endothelin1*(EDN1)* ^8, 9^.

The transcription factor ETS-related gene (ERG) facilitates binding of KLF2 to the promoter of thrombomodulin to maintain anticoagulation under LSS^10^. ERG also functions at enhancer sites of homeostatic genes^11–13^ with targets such as Jagged1 *(JAG1)*^14^, VE-cadherin or cadherin 5 (*CDH5*), claudin 5 (*CLDN5*), and von Willebrand factor (vWF). ERG balances SMAD1-3 activity to prevent endothelial-mesenchymal transition (EndMT)^15^. Loss of ERG is associated with de-repression of NFkB activity^16, 17^, and EndMT, the consequence of which is a pro-inflammatory EC phenotype associated with reduced barrier function, and de-repression of SMC proliferation. Moreover ERG is required to tether components of the SWI/SNF complex to chromatin^18^ and KLF2/4 recruits SWI/NSF to open chromatin under physiologic LSS^9^. Reduced ERG is also observed in pulmonary arteries in lung tissue from patients with PAH^19, 20^ and BMPR2 and ERG are thought to be co-regulated to maintain PAEC homeostasis^20^.

Mutations and reduced expression or function of *BMPR2* are present in patients with PAH^21^, and are also associated with EndMT^22^, characterized by an increase in the transcription factors Snail and Slug (*SNAI1, SNAI2*), a reduction in endothelial genes platelet and endothelial cell adhesion molecule 1 (*PECAM1*) and *CDH5*, and an increase in smooth muscle genes, smooth muscle alpha actin 2 (*ACTA2*) and transgelin or SM22 alpha (*TAGLN*). The mechanism whereby loss of *BMPR2* causes EndMT was related to an increase in the chromatin remodeler high mobility group antigen 1 (HMGA1)^22^ that regulates the expression of *SNAI1,2*. We have also shown that monocyte-derived endogenous retroviral protein HERV-K dUTPase can cause EndMT by inducing NFkB mediated transcription in concert with co-activators pSMAD2/3, signal transducer and activator of transcription 1 (pSTAT1) and activating transcription factor 2 (ATF2)^23^.

In this study, we exposed PAEC to the level of HSS in PAEC associated with PAH in arteries less than 500 μm^2^. We then determined whether this level of HSS mediated a reduction in BMPR2 and induced EndMT, whether the mechanism involved could be related to a reduction in KLF 2/4 or ERG mediated gene regulation, and whether restoring the transcription factor that was reduced could prevent the decrease in BMPR2 and the consequent EndMT in cultured cells. We then applied our findings to prevent PAH in a mouse with an AV shunt in which HSS induces vascular changes that cause a progressive rise in pulmonary arterial pressure and PVR.

Our studies revealed that just 24 hours of HSS applied to PAEC reduced *BMPR2* mRNA and protein and induced features of EndMT. KLF2/4 was not decreased during this time frame, but the transcriptional co-activator of KLF2/4, ERG, was reduced. Loss of ERG during LSS recapitulated EndMT and transfection of *ERG* prevented the HSS mediated reduction in BMPR2 and EndMT. Moreover, in mice with an AV shunt, delivery of *ERG* with an adeno-associated virus (AAV) targeted specifically to the lung EC attenuated EndMT and the progressive pulmonary vascular changes.

## RESULTS

### Pathologic HSS induces EndMT

We used the Ibidi flow system to deliver HSS vs. LSS to human PAEC. The PAEC were exposed to 5 dyn/cm^2^ overnight to adhere and connect with neighboring cells, followed by conditioning with LSS (15 dyn/cm^2^ for 24 h) before being either maintained in LSS for the subsequent 24 h, or switched to HSS (100 dyn/cm^2^) for 24 h (details are in the Methods section). The age, gender, race, ethnicity and cause of death of the four donor control cell lines used in these studies are in Extended Data Table S1A. We found that just 24 h of HSS vs. LSS induced features of EndMT as judged by an increase in the transcription factors, SNAI1, 2 and the mesenchymal marker ACTA2, and a reduction in EC surface markers, PECAM1 and CDH5, assessed by western immunoblot (Fig. 1A). mRNA levels were consistent with the protein levels of SNAI1, 2, PECAM1 and CDH5 (Extended Data Fig. S1A). Immunofluorescent imaging confirmed these findings and extended them by delineating an increase in another mesenchymal marker and hallmark of EndMT, fibroblast specific protein-1 (FSP1, also known as S100A4) (Fig. 1B). We next determined whether the HSS-mediated EndMT might be associated with reduced BMPR2, and indeed observed an HSS-mediated reduction in mRNA and protein levels of this receptor (Fig. 1C and D). However, the flow-induced transcription factors, KLF2/4, were not altered with HSS (Fig. 1E and F). Alignment of EC in the direction of flow and the aspect ratio of the cells increased with HSS, likely reflecting the change to a more spindle shaped mesenchymal phenotype (Extended Data Fig. S1B), and the nuclei of the EC appeared to protrude as they do in the vessels undergoing remodeling in association with HSS (Extended Data Fig. S1C).

**Fig. 1:**
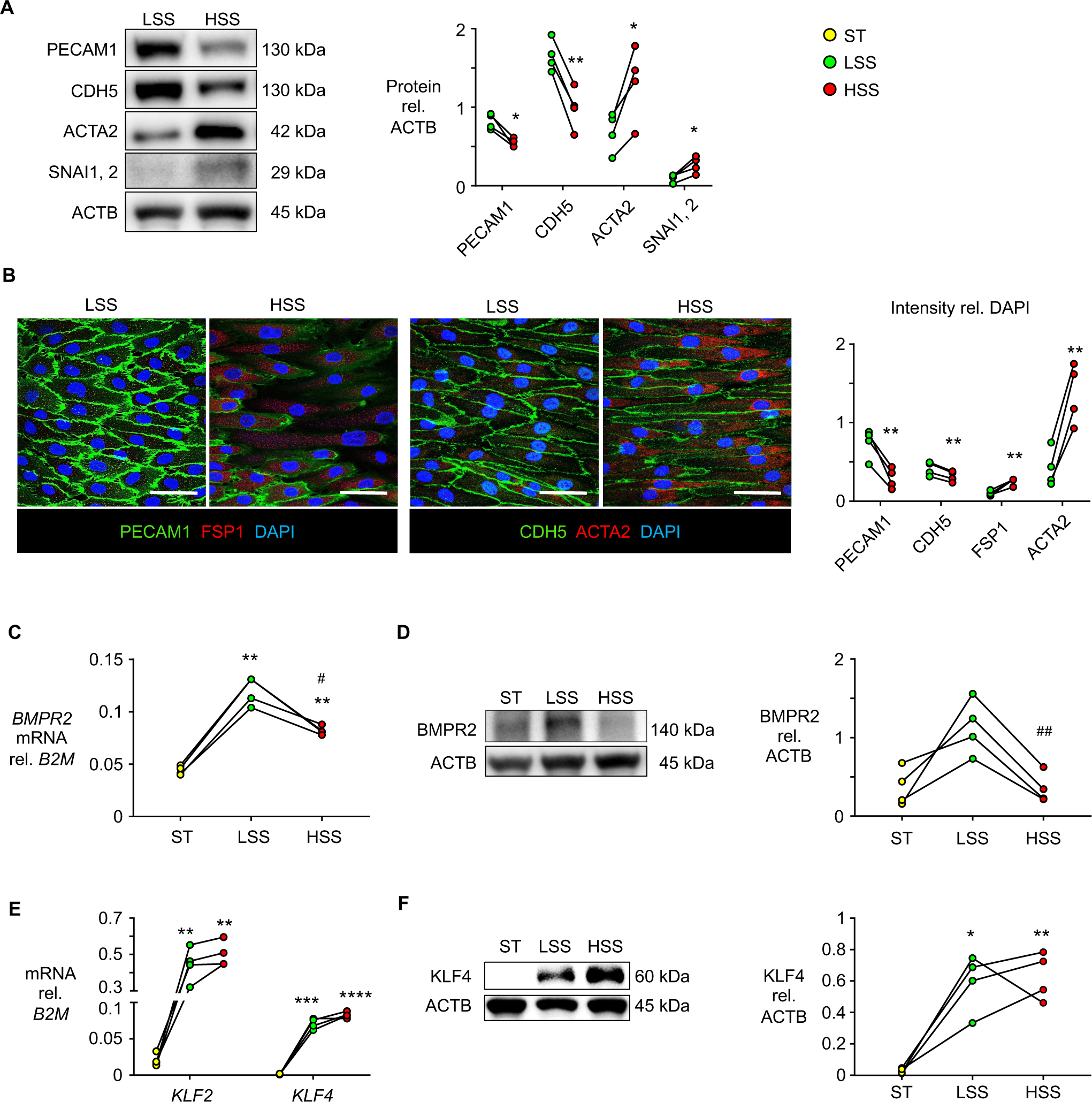
High shear stress induces Endothelial-Mesenchymal transition. Human donor PAEC exposed to high shear stress (HSS, 100 dyn/cm^2^), or laminar shear stress (LSS,15 dyn/cm^2^). Static condition (ST). **(A)** Representative immunoblot and densitometric analysis of EndMT markers relative to ACTB loading control. **(B)** Representative immunofluorescent staining of PECAM1 and CDH5 (green), FSP1 (S100A4) and ACTA2 (red), and nuclei (DAPI, blue). Quantification of intensity for each marker is shown as an average of 5 fields of view per each sample. Scale bar, 50 μm. **(C)** *BMPR2* mRNA in PAEC. **(D)** Representative immunoblot and densitometric analysis of BMPR2. **(E)** *KLF2* and *KLF4* mRNA. **(F)** Representative immunoblot and densitometric analysis of KLF4. n=4 biological replicates. In (A, B), *p<0.05; **p<0.01; ***p<0.001 vs. LSS, by paired Student t test. In (C, D, E, F) *p<0.05; **p<0.01; ***p<0.001, ****p<0.0001 vs. ST, and ^#^p<0.05; ^##^p<0.01 vs. LSS, by RM one-way ANOVA with the Geisser-Greenhouse correction, and Tukey’s multiple comparisons test.

### Pathologic HSS decreases ERG expression

To determine which transcription factor binding sites might undergo a loss of activity with HSS, we conducted H3K27ac CUT&RUN^24^ in PAEC under LSS, HSS, and static (ST) conditions.

Consistent with the increase of SNAI2 protein and mRNA levels under HSS conditions, we observed an increase in H3K27ac signal at the SNAI2 promoter, as well as in its upstream putative enhancer (Fig. 2A). Genome-wide, we identified 11,870 peaks that decrease in H3K27ac activity and 15,901 peaks that increase in H3K27ac activity under HSS conditions relative to LSS (Fig. 2B, *FDR* < 0.05). Motif enrichment analysis of the peaks that decrease in H3K27ac activity returned several candidate motifs from the SOX and ETS families of transcription factors (Extended Data Table S3). These altered motifs included ERG (Fig. 2C), a pioneer factor that can interact with KLF2^10^ and SWI/SNF^18^. Accordingly, we also observed a decrease in H3K27ac signal at peaks containing predicted ERG motifs (Fig. 2D). Proximity ligation assays further revealed that KLF4 interaction with ERG under LSS was markedly reduced with HSS (Fig. 2E).

**Fig. 2:**
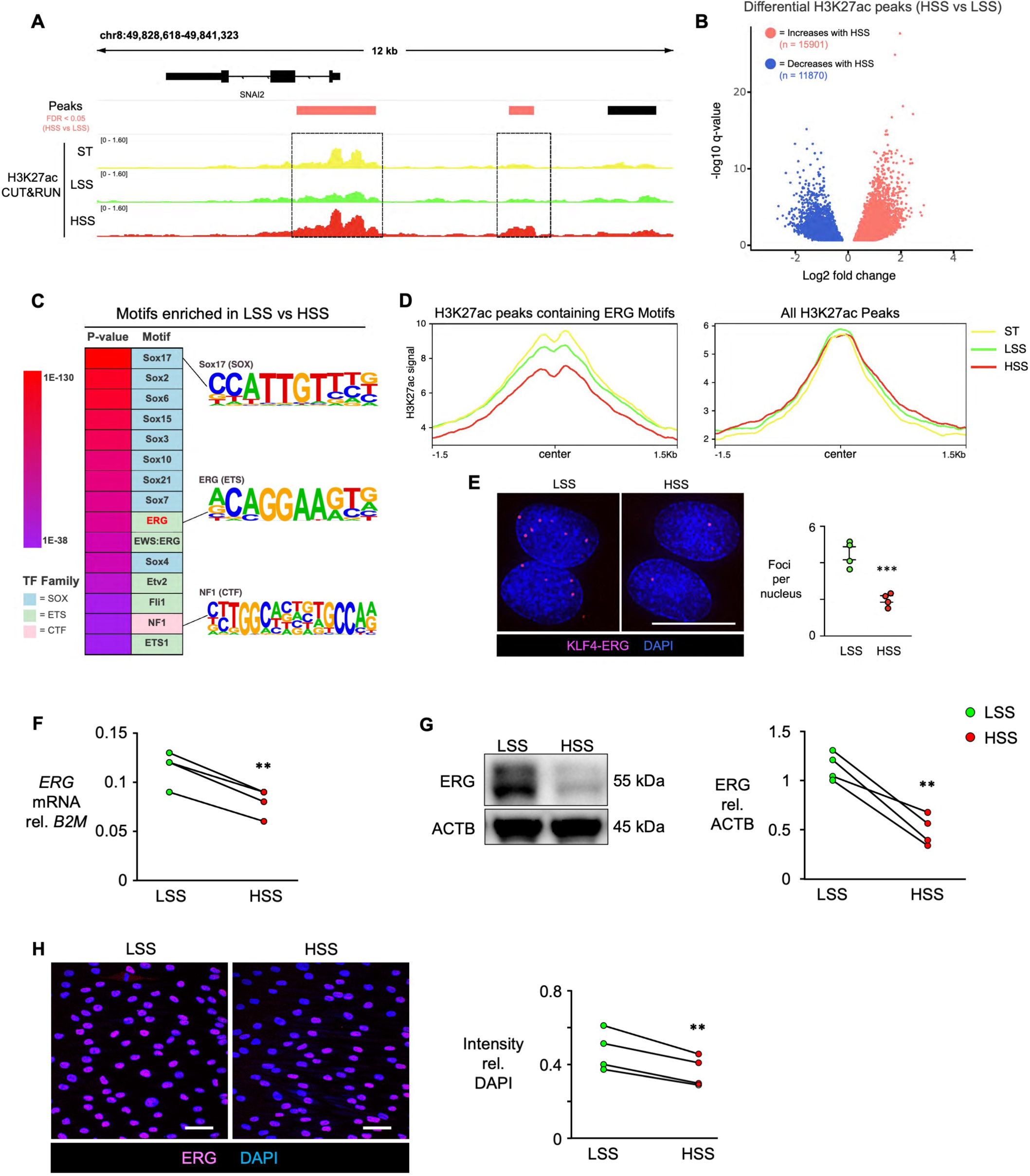
HSS reduces H3K27 acetylation at ERG motifs, and decreases ERG. Human donor PAEC cultured under ST, LSS and HSS as in Figure 1, and H3K27ac measured using CUT&RUN. (**A**) Representative H3K27ac track showing increases in H3K27 at the *SNAI2* locus under HSS. Red boxes, peaks increased in HSS vs. LSS, FDR<0.05. (**B**) Volcano plot of H3K27ac peaks with HSS vs. LSS with dots representing significant changes. P-values by a two-tailed Wald test with Benjamini-Hochberg adjustment. (**C**) Motif enrichment of peaks with increased H3K27ac in LSS vs HSS, analyzed using Homer. P-values by binomial test. Motifs colored by transcription factor family. (**D**) Normalized H3K27ac read densities for HSS, LSS, and ST across peaks containing predicted ERG motifs or across all peaks. n*=*3 experimental replicates. (**E)** Representative images of Proximity Ligation Assays of PAEC, showing interaction of KLF4 with ERG under LSS and HSS (magenta). Foci per nucleus quantified in 6 non-overlapping random fields of view per replicate (right panel). Data shown as mean±SEM, n = 4 replicates, p<0.001 by Student’s two-tailed *t*-test. Scale bar, 10 μm. **(F)** *ERG* mRNA by RT-qPCR. **(G)** Representative immunoblot and densitometric analysis of ERG. **(H)** Representative immunofluorescent staining of ERG and DAPI. Intensity quantified as an average of 5 fields of view per sample. Scale bar, 50 μm. n=4 biological replicates. *p<0.05; **p<0.01; ***p<0.001 vs. LSS by paired Student t test.

This suggested that reduced ERG may contribute to HSS-induced EndMT^15^. Indeed, HSS significantly decreased ERG mRNA and protein (Fig. 2F and 2G). These findings were reinforced by confocal imaging, where reduced nuclear staining of ERG was observed under HSS compared to LSS (Fig. 2H). We next assessed nuclear expression of ERG by confocal imaging of human lung tissue sections from PAH patients removed at transplant and in unused donor lungs as controls. Demographics and clinical data are in Extended Data Tables S1 and S2. There was a significant reduction in nuclear ERG in PAs from IPAH or APAH-CHD patients with a ventricular septal defect (VSD), when compared to control PAs (Fig. 3A).

**Fig. 3:**
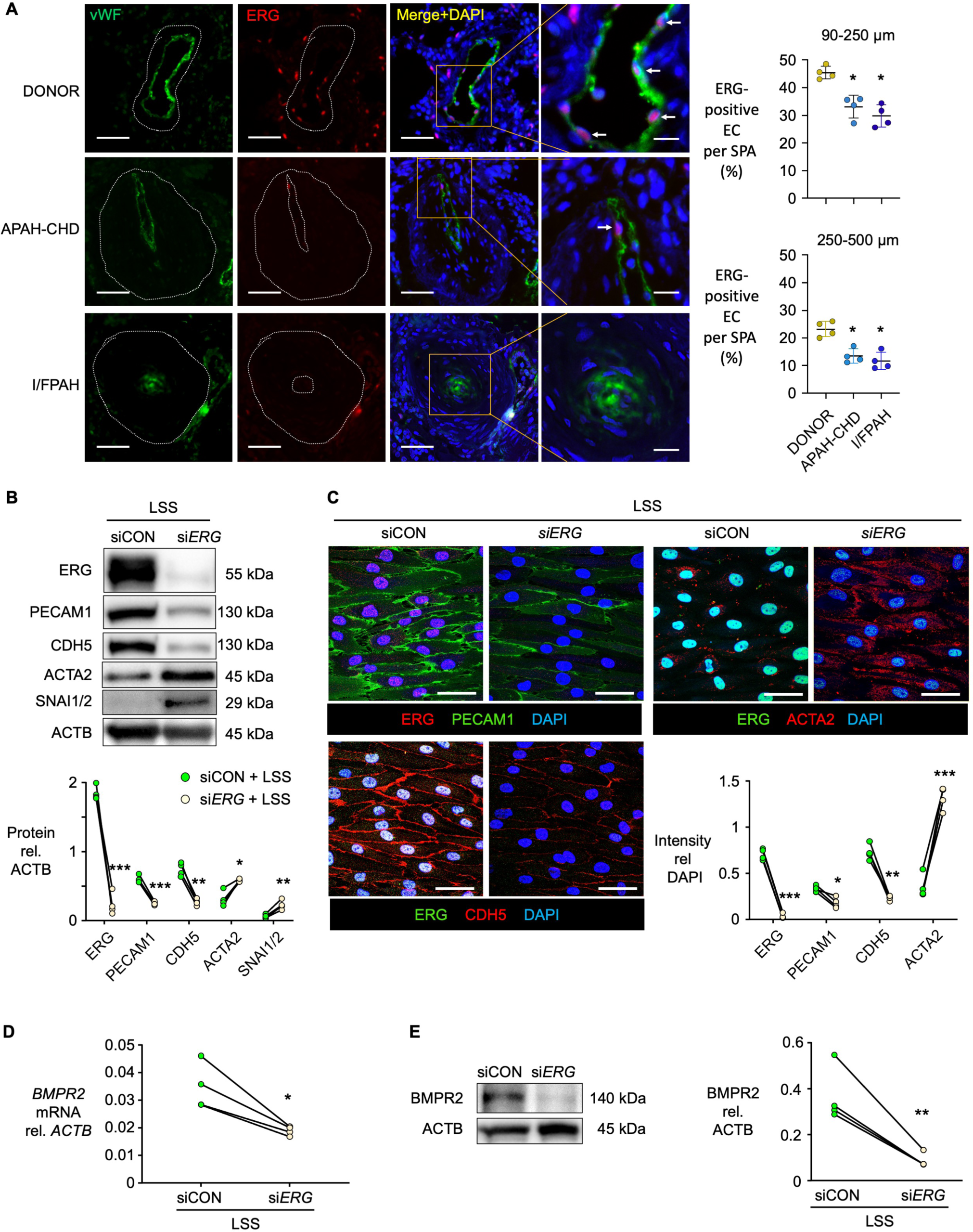
ERG is reduced in PAH; decreasing ERG under LSS recapitulates EndMT. **(A)** Confocal microscopic images of lung tissue sections from 4 patients with PAH and a congenital heart defect (APAH-CHD), 4 patients with Idiopathic or Familial PAH (I/FPAH, 2 each), or healthy controls (Donor, n=4). ERG (red), EC marker vWF (green) and nuclei stained with DAPI (blue). Arrows in magnified images (right column) point to EC with ERG in the nucleus. Scale bars, 40 μm in the three left columns, 15 μm in the magnified images. Quantification shows the average percentage of ERG-positive EC per small pulmonary artery (SPA) per patient/control, *p<0.05 vs Donor by One-way ANOVA and Kruskal-Wallis multiple comparisons test. **(B-E)** Human donor PAEC transfected with ERG siRNA (si*ERG*) or nontargeting siRNA (siCON) followed by 48 h LSS (15 dyn/cm^2^). (**B**) Representative immunoblot and densitometric analysis. (**C**) Representative immunofluorescent staining shown with intensity quantified in an average of 5 fields of view per sample. Decreasing ERG leads to reduced PECAM1 (top left), reduced CDH5 (lower left), and increased ACTA2 (Right). Scale bars, 50 μm. **(D)** *BMPR2* mRNA by qRT-PCR. **(E)** Representative immunoblot and densitometric analysis of BMPR2. n=4 biological replicates, *p<0.05; **p<0.01; ***p<0.001 vs. siCON under LSS, by paired Student t test.

### Decreasing ERG under LSS causes EndMT

To determine whether the decrease in ERG is sufficient to induce EndMT, we transfected PAEC with *ERG* siRNA, exposed the cells to LSS, and confirmed a reduction in ERG protein by western immunoblot and in nuclear expression by confocal microscopy (Fig. 3B and C) as well as a decrease in *ERG* mRNA levels (Extended Data Fig. S2A). Protein levels of PECAM1 and CDH5 were decreased and ACTA2 and SNAI1,2 were increased as assessed by western immunoblot (Fig. 3B). mRNA levels reflected reduced *PECAM1* and *CDH5* and increased A*CTA2* but *SNAI1* or *SNAI2* were not changed suggesting that the mRNA elevation of these transcription factors may be transient (Extended Data Fig. S2A). Confocal images show reduced PECAM1 and CDH5 and increased ACTA2 (Fig. 3C). *BMPR2* mRNA and protein levels were also decreased by si*ERG* under LSS (Fig. 3D and 3E). Consistent with the findings related to HSS, the aspect ratio of EC was increased with si*ERG* suggesting a mesenchymal phenotype (Extended Data Fig. S2B).

### Replacing ERG under HSS prevents EndMT

As these studies indicated that reducing ERG under LSS was sufficient to induce EndMT, we next determined whether replacing ERG under HSS would prevent EndMT. Transfection of a lentiviral *ERG* construct increased ERG under HSS and prevented EndMT as judged by increased PECAM1 and CDH5 and reduced ACTA2 and SNAI1,2 as assessed by western immunoblot (Fig. 4A). These findings were similarly observed by confocal imaging (Fig. 4B). Changes in mRNA were consistent with protein levels (Extended Data Fig. S3A). *BMPR2* gene and protein expression under HSS were also increased by transfection of *ERG* (Fig. 4C and D). The aspect ratio of the cells was reduced by expression of ERG consistent with return to an endothelial phenotype (Extended Data Fig. S3B).

**Fig. 4.**
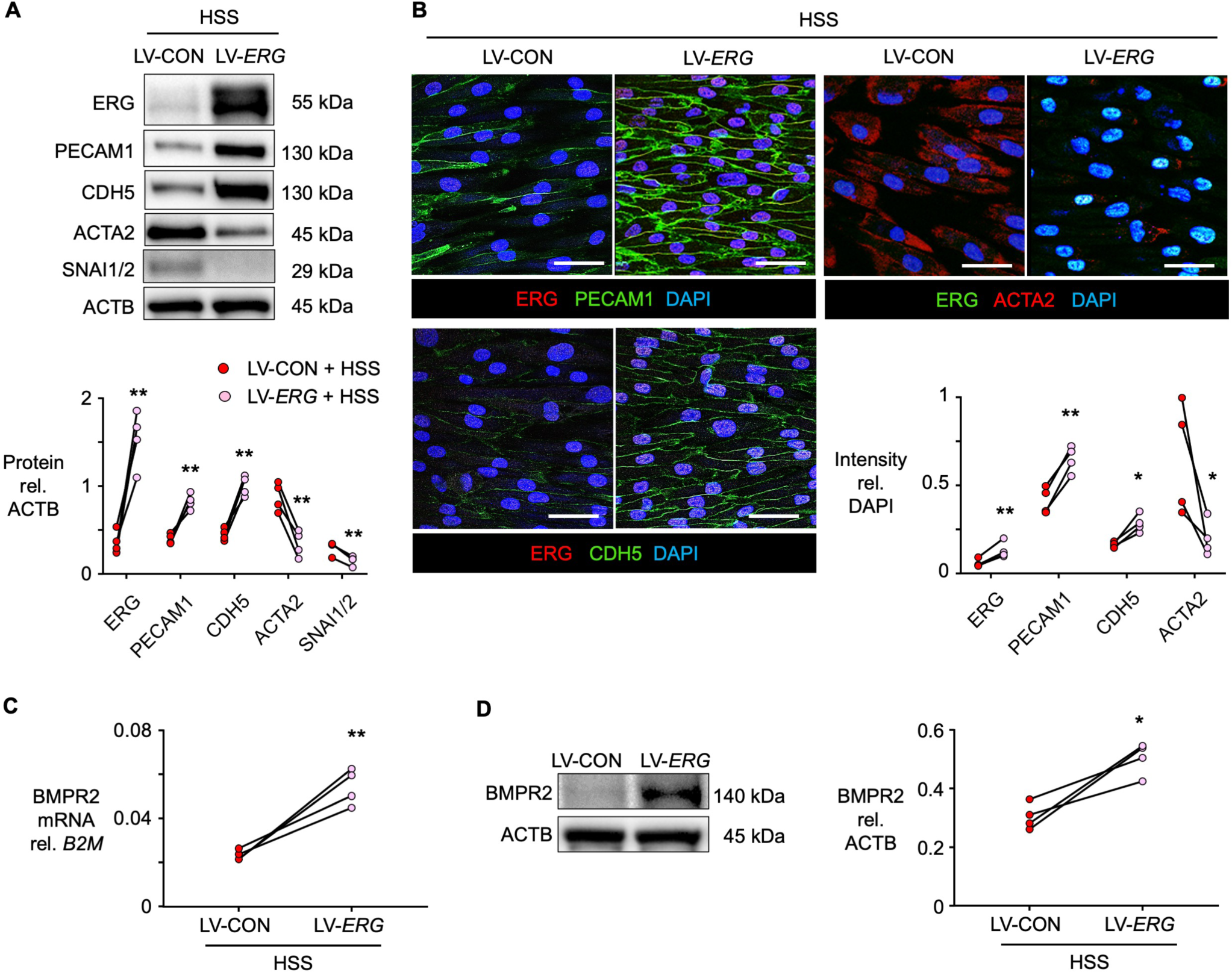
Replenishing ERG prevents EndMT under HSS. Transfecting *ERG* by lentiviral vector (LV-*ERG)* vs. control lentiviral vector carrying *EGFP* (LV-CON) in human PAEC exposed to HSS. **(A)** Representative immunoblot and densitometric analysis of ERG and EndMTmarkers, normalized to ACTB. **(B)** Representative immunofluorescent staining. Top left: Replenishing ERG (red) increases PECAM1 (green) and Bottom left: CDH5 (green). Right: Gain of ERG (green) leads to downregulation of ACTA2 (red). Quantification of intensity per nuclei stain (DAPI) shows an average of 5 fields of views per sample. Scale bar, 50 μm. **(C)** Gene expression levels of *BMPR2* in PAEC, assessed by qRT-PCR. **(D)** Representative immunoblot and densitometric analysis of BMPR2. Individual data points are shown, for n=4 biological replicates. *p<0.05; **p<0.01 vs. LV-CON under HSS, by paired Student t test.

### Pathologic HSS caused by an AV shunt induces PAH

We next related our findings to HSS produced by an AV shunt in a mouse and determined whether the progressive PAH that occurs in association with vascular changes could be related to EndMT and suppressed by targeting of ERG to lung EC. We used an AAV vector with a peptide tag previously described, AAV2-ESGHGYF, that selectively targets lung EC^25, 26^. We injected the AAV2-ESGHGYF packaged with *Erg* or a *luciferase* construct (5×10^10^ vector genomes in 100 μL PBS per mouse) one week prior to performing a sham operation or an AV shunt as previously described^27^ and as detailed in the Methods section (Fig. 5A). This AV shunt causes progressive PAH in mice when assessed up to eight weeks after placement^27^. We confirmed shunt patency by color flow Doppler examination one week after shunt placement and at the end of the study (Fig. 5A and B). Abdominal ultrasound imaging in the sham-operated mice showed a uniform color (red or blue) signal in the inferior vena cava (IVC) indicating laminar flow, whereas mice with an AV shunt showed a mosaic color pattern in an enlarged IVC, indicating disturbed flow from the abdominal aorta (Fig. 5B).

**Fig. 5.**
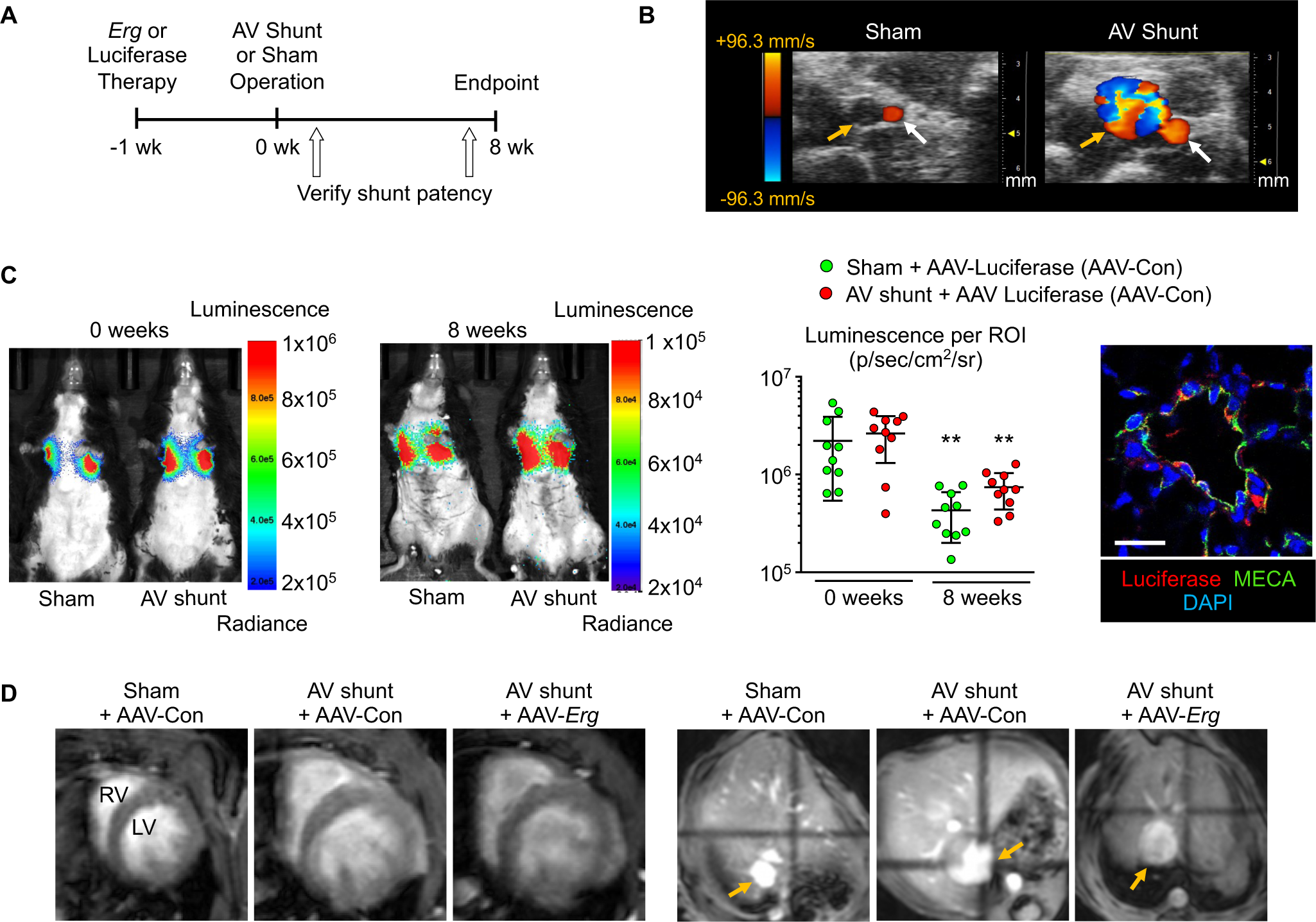
Aorto-Caval (AV) shunt in mice treated with AAV encoding *Erg* vs. Luciferase (control). **(A)** Experimental design: Mice treated with AAV encoding *Erg* (AAV-*Erg*) or luciferase (AAV-Con) one week before sham or aorto-caval (AV) shunt surgery. Shunt patency was evaluated one and eight weeks after surgery. **(B)** Patent shunt shows a mosaic Doppler color flow pattern in the IVC at the puncture site, indicating disturbed flow from aorta. Sham-operated or mice with a closed shunt show uniform color or no color at the IVC. Yellow arrow, IVC; white arrow, abdominal aorta. **(C)** Representative images of AAV-luciferase treated mice at baseline and at 8 weeks. Quantification shows luminescence Each data point represents a mouse with mean±SEM, n=10 mice per group. **p<0.01 vs. respective group at week=0, by one-way ANOVA followed by Tukey’s multiple comparisons. Right, representative image of a distal PA in a mouse treated with AAV-luciferase (AAV-Con) showing co-localization of luciferase (red) with the pan-endothelial cell marker MECA-32 (green). Scale bar, 20 μm. **(D)** Representative MRI images showing expansion of right and left ventricles (left), and expanded IVC (arrow), in the shunt groups with AAV-luciferase and AAV-*Erg* treatment vs. sham operation (right).

### Replenishing PAEC ERG prevents EndMT and attenuates PAH in AV shunted mice

To show the efficacy of targeting PAEC with the AAV vector, we monitored luciferase activity at the beginning and end of the experimental period and found that although the intensity decreased by one order of magnitude, expression was still substantial and further validated by confocal imaging of small peripheral PAs (Fig. 5C). Magnetic resonance imaging (MRI) determined that consistent with an AV shunt, dimensions of both ventricles and the IVC were increased compared to sham operation, with no impact of administration of the *Erg* construct on this feature (Fig. 5D). The increase in cardiac output (Fig. 6A) and in partial pressure of oxygen (pO_2_) in the IVC (Fig. 6B) eight weeks after the AV shunt, was also not affected by the *Erg* construct. The aortic systolic pressure was similar across all groups (Fig. 6C).

**Fig. 6.**
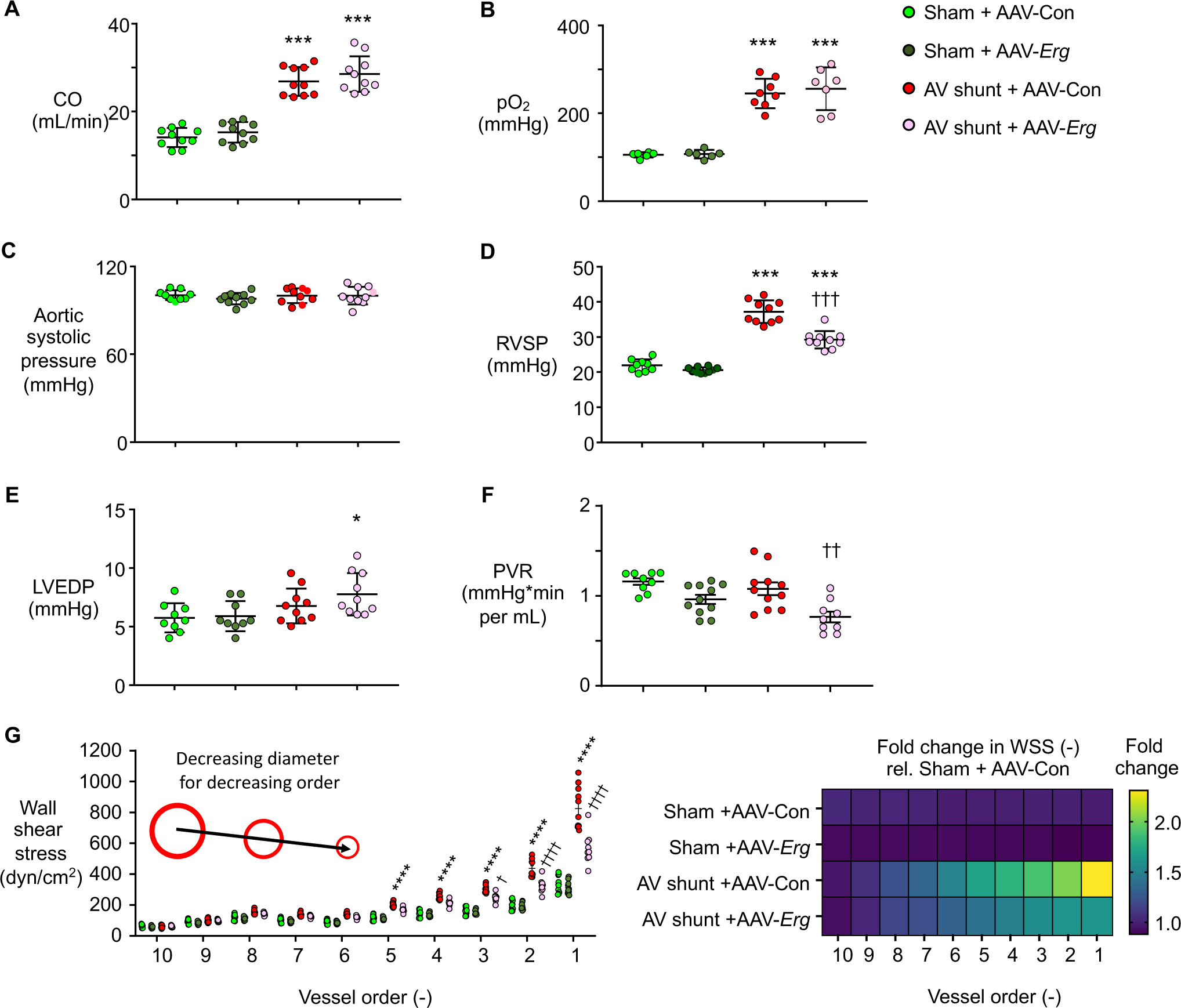
AV shunt induced PAH in mice is attenuated by replenishing PAEC ERG. Mice were treated as described in Figure 5. **(A)** Cardiac output and **(B)** partial pressure of oxygen (pO_2_) in the IVC 8 weeks following surgery indicates AV shunt patency in AAV-*Erg* and AAV-Con groups. **(C)** Aortic systolic pressure, **(D)** right ventricular systolic pressure (RVSP), and **(E)** LV end-diastolic pressure (LVEDP) determined by heart catheterization. **(F)** Pulmonary vascular resistance (PVR) calculated using catheter and echocardiography data. **(G)** Simulated values of wall shear stress (WSS) with decreasing PA vessel order generated with a reduced order computational model of blood flow (see Methods). Each data point represents a mouse with mean±SEM, n=7-10 mice per group. *p<0.05, ***p<0.001, ****p<0.0001 vs. respective Sham group; ^†^p<0.05, ^††^p<0.01, ^†††^p<0.001, ^††††^p<0.0001 vs. AV shunt with AAV-Con by one-way ANOVA followed by Tukey’s multiple comparisons.

Echocardiographic measurements performed prior to hemodynamic assessment were similar in the sham operated mice treated with the AAV-luciferase control (AAV-Con) or AAV-*Erg* vector (Extended Data Fig. S4A and B). In the AV shunt mice, left ventricular diastolic dimensions were similarly increased in AAV-Con and AAV-*Erg* groups, as was PA diameter. While stroke volume was increased, ejection fraction was similarly reduced from 60 to 50% compared to sham groups (Extended Data Fig. S4A). PAH, judged by an increased pulmonary acceleration time/ejection time, was attenuated in the AAV-*Erg* compared to the AAV-Con treated AV shunt group (Extended Data Fig. S4B). Body weight was unchanged (Extended Data Fig. S4C).

Cardiac catheterization showed that right ventricular systolic pressure (RVSP) of sham operated mice was similar with or without AAV-*Erg* treatment. Mice with an AV shunt showed PAH, as judged by elevated RVSP, compared to sham operated mice (37.2 ± 1.0 *vs.* 21.9 ± 0.6 mmHg, p<0.01), consistent with previous findings in this model at eight weeks. Treatment with AAV-*Erg* reduced the RVSP change by 50% to 29.2±0.8, (p<0.01 vs. AV shunt with AAV-Con vector) (Fig. 6D).

There was an elevation in LV end diastolic pressure (LVEDP) in the AAV-*Erg* AV shunt group when compared to sham-operated AAV-*Erg* mice but not when compared to AV shunt mice treated with the AAV-Con vector (Fig. 6E). Moreover, PVR decreased in the AV shunt mice treated with AAV-*Erg* compared to AAV-Con vector (Fig. 6F) where the expected drop in PVR with increased flow did not occur^28^ owing to vascular changes described below.

To determine the effect of replenishing ERG on the mechanobiological stimuli driving changes in EC phenotype, we used a reduced order computational model of blood flow to simulate wall shear stress (WSS) in the PA trees of each group of mice. Data from ultrasound and catheter measurements were utilized by the model to estimate the flow changes down the PA tree to allow for determination of WSS values for vessels of different sizes, where diameter decreased exponentially with vessel order. In comparing across orders for the PA tree, WSS increased in vessels of decreasing size in all groups (Fig. 6G). In PA designated orders 1-5, corresponding to vessels less than 100 μm in diameter, there was a significant increase in WSS in the AV shunt treated with the AAV-Control vector compared with the sham operated group treated with AAV-Control vector. WSS values for PA order 1-3 were lower in AV shunt mice treated with AAV-*Erg* compared to AAV-Control vector, although they were still higher than in the sham-operated mice (Fig. 6G). In PA order 4-5, WSS values in AV shunt mice treated with AAV-*Erg* were indistinguishable from controls.

Consistent with an increase in RVSP, mice with an AV shunt showed an increase in the RV weight to body weight that was reduced in those treated with AAV-*Erg* (Fig. 7A). As the LV + Septum (LV+S) to body weight ratio was similarly increased in the AV shunt mice compared with sham operated mice, the elevated RV/LV+S weight ratio in AV shunted animals treated with AAV-Con, was normal in those administered AAV-*Erg* (Fig. 7A). In sham operated mice treated with either AAV-Con or AAV-*Erg*, there was no difference in the RV or LV+S to body weight ratios, or in the weight ratio of RV/LV+S.

**Fig. 7.**
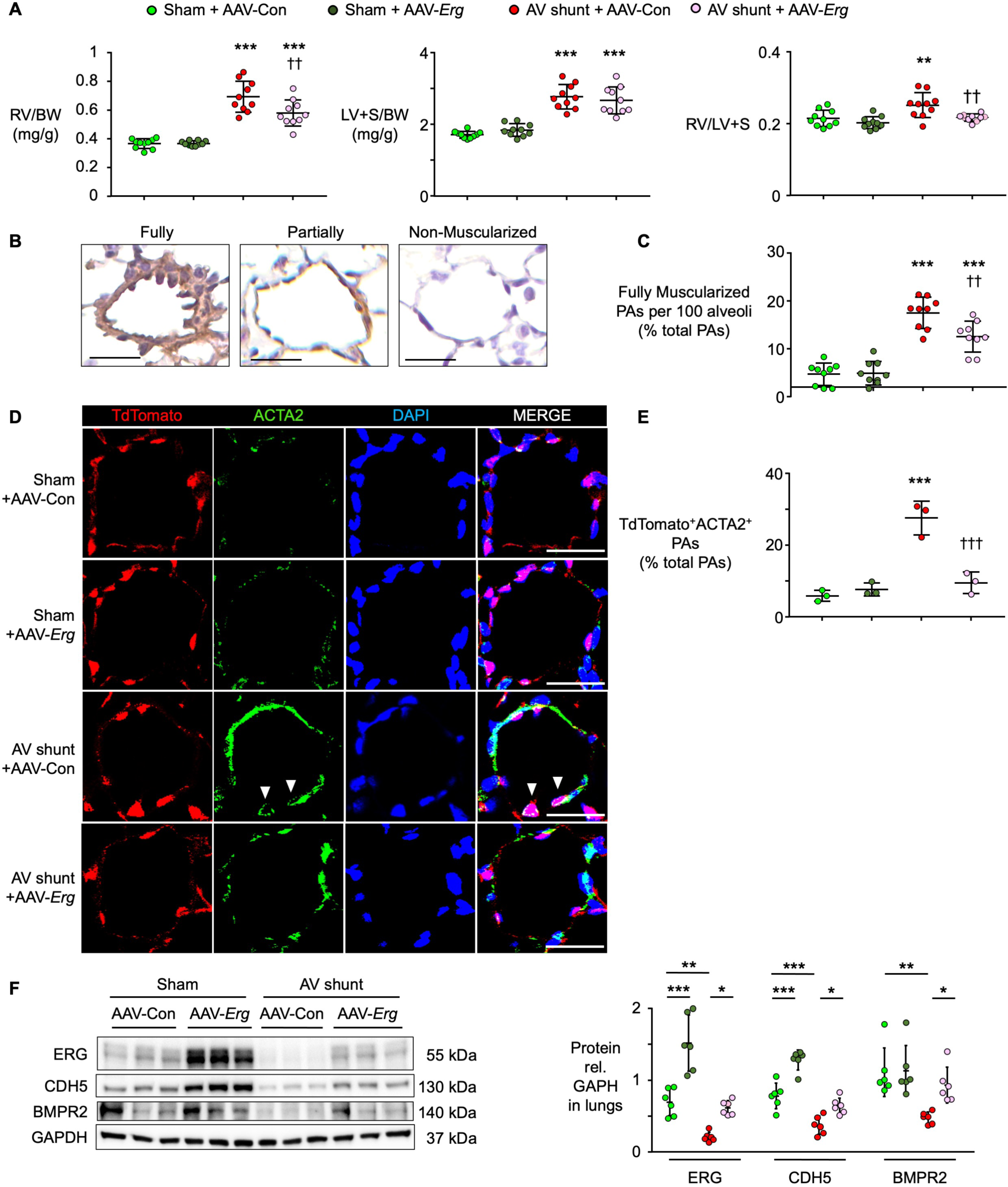
AV shunt in mice induces EndMT and remodeling of the distal PAs, which are rescued by overexpression of *Erg* in PAEC. Shunt or sham operated mice and hemodynamic parameters determined as in “Methods” and in Figure 5. **(A)** Weight ratio of the right ventricle (RV) and left ventricle (LV)+septum (S) to body weight, and the weight ratio of the RV/LV+S, eight weeks after AV shunt surgery. Each data point represents a mouse with mean±SEM, n=10 mice per group. **(B**) Examples are given of fully, paritally and non-muscular peripheral arteries. **(C)** Quantification of fully muscularized PAs per 100 alveoli in mice. Scale bars, 20 Dm. n=9 mice per group. **(D)** Immunofluorescence images show vessels with EndMT, where ACTA2 (green) is localized to tdTomato (red) positive PAEC (arrowhead), in PAs from endothelial-specific inducible tdTomato (VE-Cre/Tomato) mice with quantification of percentage of EndMT vessels in **(E)**. Scale bars, 20 Dm. An average of 30 PAs were evaluated per mouse, in n=3 mice per condition. (**F**) Representative immunoblot and densitometric analysis of ERG, CDH5, BMPR2 and GAPDH from lung tissue of mice. Mean±SEM shown for n=6 mice per group. In (**A, C, E**), *p<0.05, **p<0.01, ***p<0.001 vs. respective Sham; ^††^p<0.01, ^†††^p<0.001 vs. AV Shunt +AAV-Con. In (**F),** *p<0.05, **p<0.01, ***p<0.001 as indicated by one-way ANOVA and Tukey’s multiple comparisons test.

As expected with the elevation in RVSP, mice with an AV shunt have an increase in muscularization of distal PAs at alveolar duct and wall level as judged by ACTA2 staining (Fig. 7B and C) as well as features of EndMT judged by cells staining positive for both tdTomato (marking EC) and ACTA2. Both muscularization of distal arteries and EndMT in AV-shunt mice were reduced by treatment with AAV-*Erg* compared to AAV-Con vectors (Fig. 7D and E). There was an expected increase in ERG and CDH5 by western immunoblot in the sham operated lung of mice treated with AAV-*Erg* vector. However, the reduction in ERG, CDH5 and BMPR2 observed in the AV shunt mice treated with AAV-Con vector was abolished by treatment with AAV-*Erg* vector and values were similar to those seen in sham operated mice treated with AAV-Con vector (Fig. 7F). Figure 8 is a graphical description of our findings.

**Fig. 8.**
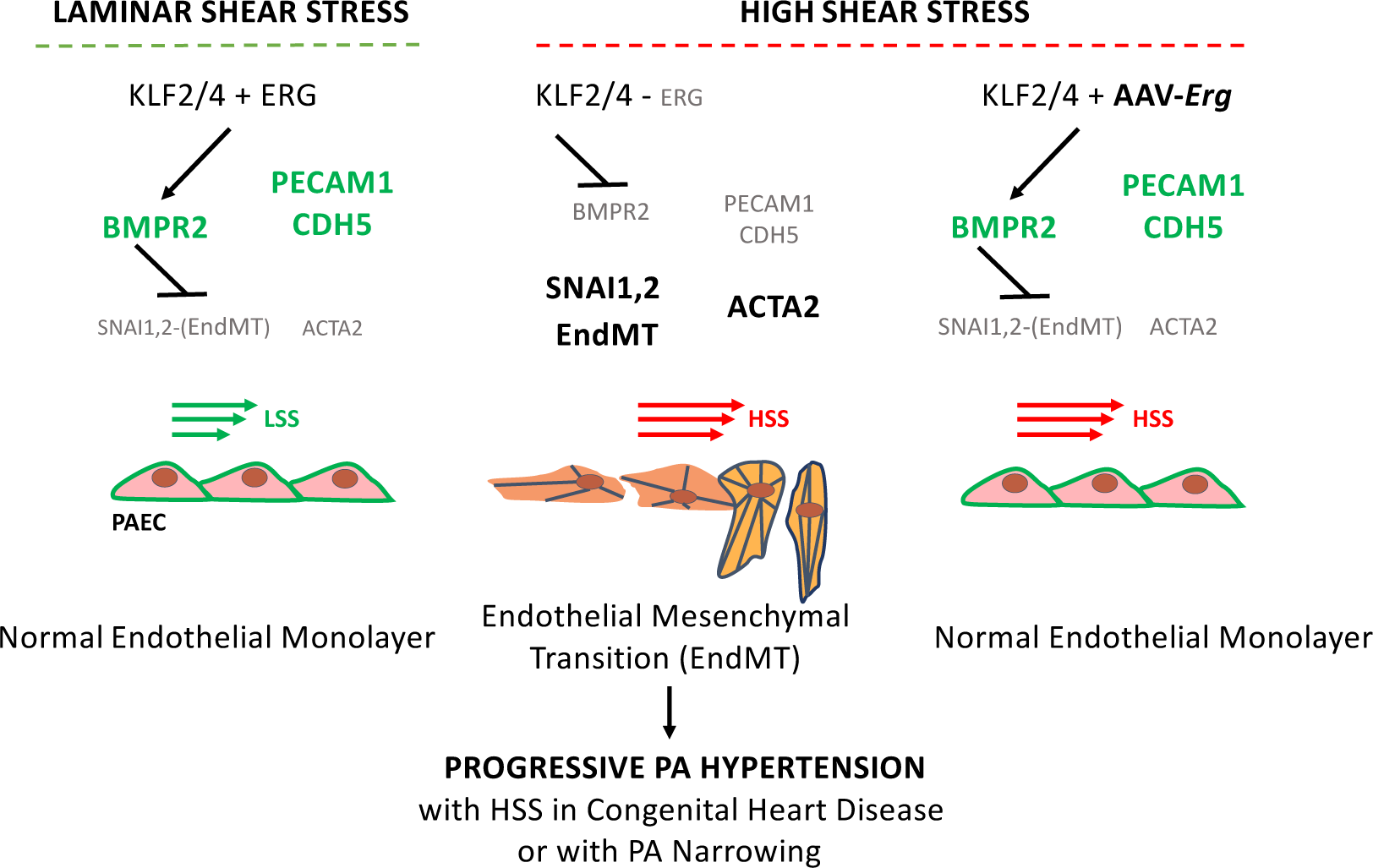
Graphical summary. HSS decreases ERG, which in turn decreases PECAM1 and CDH5, and BMPR2, inducing EndMT. Rescue of ERG suppresses EndMT under HSS.

## DISCUSSION

While some publications refer to HSS as higher than no shear stress^29^, application of pathological levels of HSS to PAEC tested in silico and applied to a cell culture system has not been studied. Computational modeling allowed us to predict the amount of pathologic shear stress to which PAEC from a child with a VSD and PAH or an individual with IPAH would be subjected, recognizing that other features, such as interaction with blood elements, SMC and the extracellular matrix, are also important in the pathophysiologic response of PAEC. In systemic EC outward remodeling has been related to proliferation and NO production under conditions of sustained high flow^30^ and transcriptomic analysis of wall shear stress above normal has been linked to increased matrix processing related to elevated mRNA levels of urokinase plasminogen activator (*PLAU*) tissue plasminogen activator (*PLAT*) tissue inhibitor of metaolloproteinase (*TIMP3*) and a disintegrin and metalloproteinase with thrombospondin motif 1 (*ADAMTS1*)^31^.

As we had previously shown that EndMT is a feature of PAH attributed to a reduction in BMPR2, our studies were designed to determine whether reduced BMPR2 is a feature of HSS. Indeed, we confirmed that BMPR2 is elevated in response to LSS but is reduced coincident with EndMT in response to HSS. As targets of KLF2/4 mediated transcription such as *BMPR2* were reduced with HSS, without a concomitant reduction in KLF2/4, we searched for enhancer/promoter motifs that were lost with HSS. Our findings showed a reduction in H3K27ac sites related to ERG binding motifs, and we subsequently showed an interaction of KLF4 with ERG that was present with LSS and reduced with HSS. Indeed, variants in ERG binding sites in enhancers marked by H3K27ac are associated with cardiovascular disease^32^, and loss of ERG is related to EndMT^33^. The reduction in ERG was pivotal to HSS induced EndMT as EndMT was recapitulated during LSS when ERG was reduced by siRNA. Moreover, transfection of *ERG* suppressed HSS mediated EndMT. Because of the cooperative nature of ERG and other ETS transcription factors, particularly FLI1^19^, it would be of interest to determine whether HSS also induces loss of FLI1 ^34^. Why ERG is reduced by HSS is not known. While impaired TEK receptor tyrosine kinase (TIE2/TEK) activation could suppress ERG^35^, we could not confirm this in PAEC under HSS. We also could not confirm a reduction in miR-96 as the cause of reduced *ERG* ^36^ under HSS. Other processes that have been related to decreased ERG include ubiquitination induced by deficiency of focal adhesion kinase^37^. Inflammatory stimuli can also degrade ERG, particularly in the lung vasculature^38^ further indicating a potential synergy between HSS and inflammation. Recently, ERG loss with aging has been implicated in the reduction in EC integrity in capillaries, a feature that predisposes to lung fibrosis^39^. Alterations in the function of the mechanosensitive Piezo1 channel might also induce the EC reduction in ERG^40^. Of interest was the observation that SOX17 motifs were reduced under HSS, given that mutations in *SOX17* and even common polymorphisms are associated with IPAH^41^ and occur in 3-7% of patients with a CHD characterized by increased PA flow and PAH^42^.

We used a capsid-modified AAV vector shown to be highly effective in targeting PAEC as a proof of concept that *Erg* delivery could attenuate HSS mediated EndMT and vascular changes. As expected, cardiac output was not altered by replenishing ERG, but PVR was reduced by delivery of the *Erg* vector, consistent with the observed reduction in RVSP and distal arterial muscularization.

While we used an AAV to deliver *Erg*, tagged nanoparticles^43^ could also be applied as delivery vectors and there may be small molecules or biologics that could be screened for their ability to increase ERG in the setting of HSS^44^. Our goal was to test the impact of maintaining ERG during HSS, but it would be interesting to determine whether increasing ERG when disease is established could induce regression of remodeling by reversing EndMT and restoring EC health and homeostasis.

## AUTHOR CONTRIBUTIONS

TS, JRM, MR conceived the project, designed the research studies, analyzed the data, and wrote the manuscript. TS, JRM, YHC, KO, LW, JMS performed cell culture and animal experiments and analyzed the data. YCL and KG analyzed the bioinformatic data related to enhancer motifs, and contributed analysis tools, under the supervision of JME. JMS carried out the computational modeling and analysis of shear stress under the supervision of ALM, and assisted with the MRI studies. JKa analyzed some of the immunofluorescent images. AC helped with statistical analyzes and data management for the manuscript. ST and SI provided technical expertise with the animal studies. MD and WY under the supervision of ALM, provided computational modeling of high shear stress for use in the cell culture studies. BDF, CP and LJP assisted with the mouse imaging studies. SS and YM supported the project and the fellowship of TS. JKö provided the AAV-peptide and related technical and scientific guidance. MR supervised the project. MR was responsible for the funding, guidance and supervision of the project.

## ACKNOWLEDGEMENTS

We thank Ms. Patricia del Rosario (Stanford University) for providing clinical information related to the cell and tissue samples, and Dr. Tushar Desai (Stanford University) for access to the Leica confocal microscope. We greatly appreciate the assistance of Dr. Michal Bental Roof in scientific and language editing, and the administrative help of Ms. Michelle Ameri. We are indebted to the Pulmonary Hypertension Breakthrough Initiative (PHBI), as the source of cells from unused donor lungs. The PHBI was funded by NIH/NHLBI R24 HL123767, the Cardiovascular Medical Research and Education Fund (CMREF) UL1RR024986. At the Stanford Center for Innovation in *In Vivo* Imaging, the Vevo 2100 system supported by NIH grant S10OD010344 and the 7T MRI scanner by NIH S10 Shared Instrumentation Grant S10RR026917-01.

## This work was supported by funding from the following sources

Japan Heart Foundation, The Mother and Child Health Foundation Kawano Masanori Memorial Public Interest Incorporated Foundation for Promotion of Pediatrics, JSPS KAKENHI Grant Number 22K15948, Pediatric Cardiology, Stanford University School of Medicine (TS); California Tobacco-Related Disease Research Program of the University of California award 27FT-0039 and Netherlands Heart Foundation award 2013T116 (JRM); National Institutes of Health grant T32 HL098049 and a Parker B. Francis Fellowship (JMS); NIH/NHLBI T32 HL129970 fellowship and Research Diversity Supplement P01 HL108797-04W1 (ST); The MSD Life Science Foundation Fellowship Grant, Japan Society for the Promotion of Science Postdoctoral Fellowship for Research Abroad and American Heart Association Postdoctoral Fellowship Award 20POST35080009 and (SI); National Science Foundation Graduate Research Fellowship DGE-1656518 (MD); Stanford Maternal and Child Health Research Institute (MCHRI) Pilot Award (ALM, MR); NIH Pathway to Independence Award K99 HG00917 and R00 HG009917, NIH-NHGRI Genomic Innovator Award R35 HG011324, The Harvard Society of Fellows and the Gordon and Betty Moore and the BASE Research Initiative at the Lucile Packard Children’s Hospital at Stanford University (JME); NIH/NHLBI grants R01 HL074186 and R01 HL 152134, and the Dwight and Vera Dunlevie Chair in Pediatric Cardiology at Stanford University (MR).

## METHODS

### Ethical Approvals

Procedures were compliant with all ethical regulations regarding animal research. Care and housing of mice was in accordance with the guidelines from the Stanford University Administrative Panel on Laboratory Animal Care (APLAC) and approved under APLAC protocol 10704. Human cells and tissues provided by the Pulmonary Hypertension Breakthrough Initiative (PHBI) were obtained under the PHBI network Institutional Review Board (IRB) protocol, with informed consent and IRB approvals at the transplant procurement sites. The cell lines and tissues were provided coded with no identifying information and therefore considered non-human subject research for the purposes of this study. The informed consent described the purpose of the lung donation and the procedure for processing the tissue.

### Cell Culture

Lungs from PAH patients and controls (unused donor lungs) were obtained though the PHBI funded by NIH (R24 HL123767) and the Cardiovascular Medical Research and Education Fund (CMREF; UL 1RR024986). Extended Data Tables S1 and S2 list demographic and clinical characteristics of the PAH patients and healthy control lungs used in the confocal imaging studies. For the cell culture experiments, we used PAEC harvested from unused donor control lungs. We used the youngest donors available to most closely resemble CHD patients; these were all male as no PAH gender bias has been shown (Extended Data Table S2). PAEC were grown in commercial EC media containing 5% FBS (ScienCell) in a 5% CO_2_ air atmosphere and used at passages 3–7. Cells were routinely tested for mycoplasma contamination. Shear stress was generated using the Ibidi Perfusion System (Ibidi). PAEC (1×10^5^ cells) were seeded in flow chamber slides (μ-Slide I 0.2 and 0.4 mm ibiTreat; Ibidi) and grown to confluence. Cells were first exposed to 5 dyn/cm^2^ overnight, then to 15 dyn/cm^2^ of unidirectional uniform LSS for 24 h and either exposed to an additional 24 h of LSS, or 24 h of 100 dyn/cm^2^ unidirectional uniform HSS. Static (ST) controls were performed simultaneously with the shear stress experiments, cultured on 0.4 mm chamber slides using the same EC media. Medium was changed every 24 h, to avoid nutrient deprivation that can occur in flow chamber slides that are not perfused.

### CUT&RUN

CUT&RUN was performed using the CUTANA CUT&RUN kit (#14-1048, EpiCypher, USA) following manufacturer’s instructions. Briefly, 100,000 cells were detached using Accutase (Stem Cell Technologies, USA) and centrifuged at 600 x g for 3 min. The pellet was resuspended in 100 μL wash buffer and centrifuged for 3 min at 600 x g. This was repeated twice. After resuspension in 100 μL wash buffer, the cells were incubated with 10 μL of activated Concanavalin A conjugated paramagnetic beads on a rotator for 10 min at RT. Tubes were placed on a magnetic stand and the supernatant was carefully removed. The pellet was resuspended in 50 μL cold Antibody buffer with 0.5 μg of antibodies targeting H3K27ac (ab4729, Abcam, USA), or H3K27me3 antibodies and Rabbit IgG provided with the kit as positive and negative controls, respectively. Samples were incubated on a nutator overnight at 4°C. The next day, the tubes were placed on a magnetic stand and the supernatant was removed, followed by resuspension, and washing with cold cell permeabilization buffer. The pellets were resuspended with 50 μL cell permeabilization buffer with 2.5 μL/reaction of pAG-MNase, and allowed to incubate for 10 min at RT. Following removal of the supernatant in a magnetic stand, the pellet was washed twice with 200 μL of cold cell permeabilization buffer and resuspended in 50 μL cell permeabilization buffer. The tubes were placed on ice and 1 μL/reaction of 100 mM calcium chloride was added to activate digestion. Tubes were incubated on a nutator for 2 h at 4°C. The reaction was terminated by adding 33 μL Stop Master Mix comprised of stop buffer and 1 μL of E. coli spike-in DNA and samples were placed in a thermocycler and incubated at 37°C for 10 min. Following a quick spin, the tubes were placed on a magnetic stand and the supernatant was transferred to fresh tubes. DNA was purified by adding 420 μL of DNA binding buffer to each reaction and loading into DNA Cleanup Column + collection tubes provided with the kit. Columns were centrifuged at 16,000 x g for 30 sec at RT, washed twice with 200 μL DNA wash buffer, and finally DNA was eluted using 12 μL of DNA elution buffer. Paired-end sequencing libraries were prepared using the CUTANA™ CUT&RUN Library Prep Kit (#14-1001, EpiCypher, USA) following manufacturer’s instructions. Enrichment of mononucleosomal fragments was confirmed and library concentration was determined using Agilent Bioanalyzer.

### Data Analysis

Data were pre-processed using the ENCODE ATAC-seq pipeline (https://github.com/ENCODE-DCC/atac-seq-pipeline). Briefly, this pipeline takes raw FASTQ files as input, trims adaptors with Cutadapt 1.9.1, aligns reads with Bowtie2, and filters unmapped or duplicated reads with Samtools 1.7 and Picard 1.126, outputting filtered, deduplicated reads aligned to the hg19 reference genome. Peaks for each condition were then called using MACS2. Reads in peaks were counted using DiffBind 2.4.8 (https://rdrr.io/bioc/DiffBind/) to produce a count matrix. Differentially bound peaks were detected using DESeq2 (https://bioconductor.org/ <u>packages/release/bioc/html/DESeq2.html</u>) with an FDR cutoff of <0.05. Data was visualized using the Integrative Genomics Viewer (http://www.broadinstitute.org/igv). The HOMER (http://homer.salk.edu/homer/) function findMotifsGenome was used with default parameters to search for motif enrichment in peaks with differential H3K27ac.

### siRNA Transfection

PAEC were transfected with ON-TARGETplus SMARTpool RNAi targeting *ERG* (L-003886-00-0005) (siERG) or ON-TARGETplus non-targeting pool (D-001810-10-05, Dharmacon) as siControl (siCON). Transfection was performed using 20nM of siRNA with Lipofectamine RNAiMAX (Lifetechnologies) in Opti-MEM 1 reduced serum medium (ThermoFisher) for 6 h, after which media were changed to regular ECM (ScienCell). Transfected PAEC were seeded in flow chamber slides 24 h after the start of transfection, incubated for 12 h, and 36 h after the start of transfection, the PAEC were exposed to LSS for 48 h.

### Cell Transfection

Lentiviral vector encoding human *ERG* (pLV[Exp]-mCherry:T2A:Puro-CMV>hERG [NM_001243428.1] (Vector ID: VB200817-1810bfd or eGFP control vector (pLV[Exp]-Puro-CMV>EGFP (VectorBuilder VB900088-2219pdm) were obtained from VectorBuilder. The pLV vectors were co-transfected with packaging plasmids (Celecta #CPCP-K2A) into HEK 293T cells using Lipofectamine 3000 reagent (Invitrogen #L3000001) according to the manufacturer’s recommendations. Media was changed to 5% FBS ECM media 24 h after transfection and the viral supernatant harvested at 48 and 60 h. For infection, viral supernatants were added to human PAEC together with 8 μg/mL polybrene (Santa Cruz #sc-134220) immediately after each harvest. Media were exchanged to fresh ECM media at 72 h. Infected cells were selected with puromycin (1 μg/mL). Transduction efficiency was confirmed by immunoblotting.

### Immunohistochemistry (IHC)

Mouse sections from paraffin-embedded lung tissues were de-paraffinized and rehydrated. Epitope retrieval was performed by boiling the sections in citrate buffer (pH 6.0, Sigma Aldrich #C999) for 20 min. Sections were incubated with 0.3% hydrogen peroxide in methanol for 30 min to block endogenous peroxidase, washed, and blocked with 5% BSA+0.2% Triton X-100 in TBS (TBST) for 1 h at room temperature. The sections were incubated with rabbit primary antibody for ACTA2 (Abcam # ab124964) using Dako animal research kit (Dako #A3954) per manufacturer’s instructions. Sstained sections were developed with DAB+ substrate-chromogen for 2 min and counterstained with hematoxylin. Images were acquired using a BZ-X710 microscope (Keyence) with 20x objectives. Muscularization was determined by ACTA2 staining of the vessels and assessed by comparing the number of muscularized vessels at alveolar duct and wall to the total number of distal small vessels at this level (diameter <50 μm). The number of these small pulmonary vessels per 100 alveoli was assessed by comparing the number of vessels to the number of alveoli in same image. Antibodies used in the immunohistochemistry studies are listed in Extended Data Table S4.

### Immunofluorescence

#### Immunofluorescence of PAEC

PAEC were cultured on flow slides (μ-Slide I 0.2 or 0.4 mm ibiTreat; Ibidi), washed with PBS and fixed with 4% PFA for 15 min. After washing with PBS, slides were blocked with 5% normal donkey serum (EMD Millipore Corp) and 0.5% Triton X-100 (Sigma Aldrich) in PBS at room temperature for 1 h prior to incubation with primary antibodies listed below at 4°C overnight. Secondary antibody incubations were performed using 1:400 dilutions (all from ThermoFisher Scientific), in the blocking buffer at room temperature for 1 h. Slides were incubated with DAPI (4,6-diamidino-2-phenylindole) (ThermoFisher Scientific). Stained slides were imaged using Leica Application Suite X software on a Leica Sp8 (Leica). Quantification of the nuclear fluorescence intensities was performed using ImageJ.

Paraffin-embedded tissue sections (7 μm thick) obtained from lungs removed at transplant from PAH patients (4 APAH-CHD with a ventricular septal defect, 2 FPAH with a *BMPR2* mutation and 2 IPAH) or from 4 unused donor lungs (donor controls) were deparaffined and rehydrated. Antigen retrieval was performed by heating the slides in 1x Citrate buffer pH 6 (Sigma) for 40 min. Sections were blocked and permeabilized in blocking buffer (3% BSA plus 5% goat, 0.2% Triton X-100 in 1XTBS) for 1 h at RT followed by incubation with primary antibodies against ERG (Santa Cruz, 1:100) for 72 h and EC marker, vWF (Abcam, 1:250) for 48 h at 4°C. Sections were washed three times with 1X PBS followed by incubation with IgG-conjugated secondary antibodies (goat anti-mouse IgG overnight and Alexa Fluor™ 488 goat anti-rabbit IgG for 4 h (both from ThermoFisher Scientific, 1:1000). Sections were washed three times as above and followed by 1 h incubation with Alexa Fluor™594 Tyramide SuperBoost for ERG staining. Secondary antibody only controls were used to assess background and selectivity of staining. Nuclei were stained with DAPI. Images were acquired by a blinded observer using a Leica TCS SP8 confocal microscope (JH Technologies, San Jose, CA). Images were taken from immunostaining done at the same time and under the same conditions for a given experiment, and images shown best-represent the average level of each marker for each condition studied. Positive staining on lung sections was based on colocalization with an EC marker and the location of the protein of interest was noted as nuclear.

#### Immunofluorescence imaging of mouse tissues

Paraffin-embedded mouse lung tissue sections were deparaffined and rehydrated. We did not boil the sections for epitope retrieval, as boiling abolished the tdTomato signal. Sections were reacted with 0.3% hydrogen peroxide to block endogenous peroxidase, and with 1% BSA in PBS to block non-specific staining^23^. Then, the sections were incubated with primary antibodies against the protein of interest overnight at 4°C. Following overnight incubation, sections were washed three times with PBS and incubated with Alexa 488 (labeled anti-mouse or rat, 1:400) or Alexa 594 (labeled anti-rabbit, 1:400)-conjugated secondary antibody (Invitrogen, Waltham, MA) for 1 h at room temperature. Sections were washed three times with PBS, mounted with mounting medium containing DAPI (Vector Laboratories, Burlingame, CA) and stored at 4°C until analysis. Confocal analysis was performed using a confocal laser-scanning microscope (FV1000, Olympus, Center Valley, PA).

Antibodies used for immunofluorescence are listed in Extended Data Table S4.

### Western Immunoblot

Cultured cells on a μ-Slide were washed with ice-cold PBS, and lysates prepared by adding 150μl of Piece RIPA Buffer (Thermo Scientific #89901) with Halt protease and phosphatase inhibitor (Thermo Scientific #78442) on ice. Cell extracts were collected using Amicon Ultra-0.5 (Sigma-Aldrich) at 4°C. Protein concentration was determined by the BCA assay (Thermo Scientific #23228). Equal amounts of protein were mixed with sample buffer (Thermo Fisher Scientific NuPAGE LDS Sample Buffer #NP0007) containing TCEP (Pierce #77720) and were separated by SDS-PAGE on Bolt 4%–12% Bis-Tris Plus gels (Thermo Scientific #NW04122) and transferred onto polyvinylidene difluoride (PVDF) membranes (Biorad #1620177) at 120V for 90 min at 4°C. PVDF membranes with proteins were washed with 0.1% Tween-20 in Tris-buffered saline (TBST) for 5 min and blocked with TBST + 5% bovine serum albumin (BSA; Sigma Aldrich #A3059) at room temperature for 1 h. Next, the membrane was incubated in 5% BSA-TBST with primary antibodies overnight at 4°C. After overnight incubation, the membrane was washed three times with TBST for 15 min and incubated with secondary horseradish peroxidase (HRP) antibody (1:5,000) for 1 h at room temperature. Membranes were developed using chemiluminescence reagent Clarity Western ECL Substrate (BioRad #1705061) or SuperSignal West Femto Maximum Sensitivity Substrate (Thermo Scientific #34096) with BioRad ChemiDoc XRS system. Densitometric quantifications were performed using ImageJ. Extended Data Table S4 lists the details of the antibodies used for Immunoblotting.

### Reverse-Transcriptase qPCR (RT-qPCR)

Total RNA was extracted and purified from cells using the Quick-RNA MicroPrep Kit (Zymo Research #R1051). The quantity and quality of RNA was determined using a Synergy H1 Hybrid Reader (BioTek) and then RNA was reverse transcribed using the High-Capacity RNA to cDNA Kit (Applied Biosystems #4387406) according to the manufacturer’s instructions. RT-qPCR was performed using 1 μL of 5 μM mixed primers, 5 μL Power SYBR green PCR Master Mix (Applied Biosystems # 4309155), 2 μl dH_2_O and 2 μL cDNA sample. The qPCR program consisted of 50°C for 2 min, 95°C for 30 sec, followed by 40 cycles of 95°C for 15 sec, and 60°C for 60 sec. Each measurement was carried out in duplicate or triplicate using a CFX384 Real-Time System (Bio-Rad) in a 10 μL reaction and performed according to the manufacturer’s instructions. Primer sequences were designed using NCBI’s Primer-BLAST function or PrimerBank (http://pga.mgh.harvard.edu). Data were analyzed by the delta-delta CT method. Expression levels of selected genes were normalized to β-Actin or β2M. Extended Data Table S5 lists the primer sequences used for RT-qPCR.

### Proximity Ligation Assays (PLA)

PLA in cultured PAEC were performed using Duolink PLA protein detection technology (Millipore Sigma) according to manufacturer’s instructions. Briefly, slides with PAEC were crosslinked with 4% PFA for 10 min, washed and incubated with Duolink Blocking solution for 1 h at 37°C. Slides were incubated with rabbit antibodies targeting KLF4 (1:100, sc20691, Santa Cruz Biotechnology) and mouse antibodies targeting ERG (1:100, sc376293, Santa Cruz Biotechnology) overnight at 4°C. Rabbit and mouse IgG were used as controls. The next day, slides were washed, and then incubated with the Duolink PLUS and MINUS probes for 1 h at 37°C. After washing the slides, the probes were ligated for 30 min at 37°C. Slides were washed and rolling circle amplification was performed for 100 min at 37°C, after which slides underwent final washes and were mounted using Duolink in situ mounting medium with DAPI (Millipore Sigma). Stained slides were imaged using Olympus Fluoview FV1000 software on an Olympus FV1000 confocal laser scanning microscope (Olympus). Quantification of the average number of foci per nucleus was performed using ImageJ.

### Mouse model for EndMT

To monitor EndMT, mice with an endothelial-specific inducible tdTomato were created by breeding VE-cadherin-CreER and tdTomato^fl/fl^ mice (VE-cadherin-CreER/tdTomato) in our laboratory. The tdTomato transgene was expressed in EC by administering 2 mg tamoxifen per day intraperitoneally for 8 days, to mice 6 weeks of age. Male transgenic mice were randomly assigned to treatment or control group with or without AV shunt^23^. We used only male mice because their larger size facilitated shunt placement and because CHD-APAH occurs in both female and male patients with no sex differences,

### Aorto-Caval Arteriovenous (AV) Shunt in Mice

Aorto-caval arteriovenous (AV) shunt to induce pulmonary remodeling was established by an abdominal aorta-inferior vena cava shunt operation using a 25G needle to puncture both walls of the aorta into the interior vena cava (IVC) at two sites as described by Okamura et al ^27^. Briefly, C57BL/6J male mice at 8 weeks of age were anesthetized with an inhalation of 1-2% isoflurane in Oxygen (1 L/min). After removal of the fur, a midline abdominal incision was made, and the abdominal aorta between renal artery and common iliac artery dissected for clamp and puncture placements. After clamping the aorta at a distal site in a branch of the renal arteries, a 25G needle was used to puncture the aorta through IVC to create the AV shunt. Hemostatic compression was applied immediately after the removal of the clamps, using sterile cotton swabs. Nylon monofilament Prolene 4/0 sutures were used to close the peritoneum and skin. Sham operated mice were treated by the same experimental protocol, except for the shunt procedure. Successful creation of the AV shunt resulted in pulsatile arterial blood flow in the IVC.

### AAV Administration to Mice

The lung EC targeting AAV2-ESGHGYF vector from the Körbelin Lab (University Medical Center Hamburg-Eppendorf, Germany)^25^ was used for all AAV experiments. Recombinant AAV2-ESGHGYF particles carrying mouse *Erg* (NCBI RefSeq NM_133659, Vectorbuilder vector ID: VB210511-1276jbk) or luciferase as a control (VectorBuilder vector ID: VB210511-1278xwa) were obtained from VectorBuilder. The *Erg* or luciferase AAV vectors were injected in the tail vein at a dose of 5 × 10^10^ vector genomes (vg) in 100 μL PBS per mouse. One and nine weeks after administration, animals injected with luciferase vector were anesthetized with isoflurane (1% in 1L O_2_/min) and Luciferin substrate (Biosynth L-8220) was injected into the intraperitoneal cavity of mice at 150 mg/kg. Bioluminescence was visualized 20 min after injection with the LagoX (Spectral Instruments Imaging) *in vivo* imaging tool.

### Abdominal Ultrasound

Mice undergoing the AV shunt operation were evaluated for shunt patency one and eight weeks after the surgery by color Doppler ultrasound, using the MS550S probe in the Vevo 2100 system (FUJIFILM Visual Sonics, Toronto, ON, Canada). Shunt patency is indicated by a mosaic pattern in the IVC at the puncture site. Mice where the shunts had closed were excluded from the study.

### Cardiac Magnetic Resonance Imaging (MRI)

To visualize changes in the heart and inferior vena cava, representative animals from the Sham + AAV-Con, AV Shunt + AAV-Con, and AV Shunt + AAV-*Erg* groups were scanned on a Bruker 7T MRI (Bruker Corp., Billerica, MA) within 48 hours of sacrifice for RVSP assessment. Mice were anesthetized with isoflurane (1.5 - 2.5% in 1 L O_2_/min) and given Gadavist (gadobutrol; Bayer) at a dosage of 0.1 mmol/kg via tail vein injection immediately prior to imaging to enhance contrast. Three axial slices were acquired at the level of the papillary muscles (TR: 50 ms, TE: 2.6 ms, Flip angle: 45 deg, Field of View: 25 mm, Image matrix 256 x 256, Thickness: 0.8 mm) and the inferior vena cava below the diaphragm (TR: 42 ms, TE: 2.5 ms, Thickness: 1.0 mm).

### Two-Dimensional Echocardiographic Assessments

One to two days before right ventricular systolic pressure (RVSP) assessment, two-dimensional echocardiographic measurements of cardiac function were carried out using the MS550S probe in the Vevo 2100 system in unventilated mice under isoflurane anesthesia (1.5% - 2.5% in 2 L O_2_/min). Echocardiographic images were used to determine the left ventricle (LV) end diastolic and systolic diameters, the aortic diameter (AOD) - close to the LV outflow tract, heart rate (HR), the velocity time integral (VTI) of the LV outflow tract, PA diameter and the pulmonary arterial acceleration time (PAAT). The percentage of LV fractional shortening (% LVFS) and the percentage of the LV ejection fraction (% LVEF) were calculated as described by Stypmann et al^45^. Left ventricular cardiac output (LVCO) was calculated as: Stroke Volume x HR = [(AOD/2 x AOD/2) x 3.1416] x VTI x HR. Mice were monitored during the echocardiographic assessment, as LVCO is distinctly dependent on heart rate.

### Hemodynamic Assessments

Cardiac catheter measurements of RVSP were carried out under isoflurane anesthesia (1.5% - 2.5% in 2 L O_2_/min) in unventilated mice, using a 1.4 Fr Millar catheter (Millar Instrument, Houston TX, Model SPR-671) through the jugular vein, and recorded with the PowerLab 7 system (AD Instruments PowerLab 7, Colorado Springs CO) as previously described^46^. Cardiac catheter measurements of the left ventricular end-diastolic pressure (LVEDP) were obtained via the carotid artery under closed chest technique^47^.

### Exclusion Criteria for the Mouse Experiments

Mice were excluded from the defined experimental endpoint if they displayed alopecia, hunched back, reduced body weight for no reason, or body weight that was significantly higher than normal range for their age, or showed abnormal behavior during the experimental period. Mice were euthanized prior to the defined experimental endpoint, and not included in the analysis if they displayed any of the following signs: Inability to ambulate that prevents the animal’s easy access to food and/or water; Inability to maintain itself in an upright position; Prolonged (greater than 48 h) in appetence and/or clinical dehydration; weight loss of 10-15% within a few days, or overall weight loss of 20%; Agonal breathing and cyanosis; chronic diarrhea or constipation; were unconsciousness with no response to external stimuli such as a toe-pinch withdrawal test. Mice that died due to bleeding caused by the RVSP and LVEDP procedures were excluded. In accordance with these criteria, we excluded mice designed for the animal experiment due to failed RVSP and LVEDP measurement.

### Method for Blinding of the Mouse Experiments

We utilize capital letters as a code to identify transgenic mice colonies and a numerical sequence for the cage IDs to avoid repetition. Up to five mice were weaned into each cage. An ear notch was created and defined as a number at the time of collecting tails for genotyping (left top = 1, left middle = 2, left low = 3, right top = 4, right middle = 5). The identification of the mice was based on the cage number followed by the ear notch number with the colony code. We thus generated an individual mouse ID that was unique and did not disclose the genotyping results. An additional ear notch on the right ear (right low = 6) was introduced when alternating mice from different cages/groups to avoid repetition.

### Computational simulation of wall shear stress (WSS) in the PA tree

Pulmonary arterial tree hemodynamics were simulated with Strahler diameter-defined morphometric trees, as previously used in human modeling and extended for murine applications^1, 2, 48^. This provided a reduced order modeling framework for estimation of wall shear stress in vessels throughout the PA tree, that was adapted in this work for murine modeling using collected data and assumptions based on the literature^49, 50^. Vessels were organized into *N* total orders with diameter and length decreasing with decreasing order *n*. As there is strong evidence across multiple species that diameter and length of PAs tend to decrease with linear scaling factors *ϕ_d_* and *ϕ_l_* from order-to-order, diameters *d_n_* = (*ϕ_d_*)*^n^d_N_* and lengths *l_n_* = (*ϕ_l_*)*^n^l_N_* were used to assign the size of vessels down the tree, where *d_N_* and *l_N_* are the diameter and length of the highest order vessel, respectively. A connectivity matrix *C_mn_* was prescribed that defines the average number of vessels of order *m* connected to each other order *n*. The PA tree structure was then generated from the top down by iteratively adding vessels to the tree according to previously defined rules^2, 49^, where each element is subdivided and outlets are added based on the total from the appropriate column of the connectivity matrix to create a series of asymmetric bifurcating junctions. Hemodynamics in the tree were then solved with the assumption of Poiseuille flow within each segment, and wall shear stress *τ_v_* in each vessel segment *v* was calculated according to 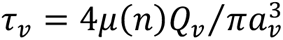 with viscosity *μ*(*n*) varying for each order according to the Fahraeus-Lindqvist effect, volumetric flow in each vessel *Q_v_* determined from splitting the flow of each parent vessel proportionally according to the distal resistances of its daughters, and radius *a*_*v*_ determined from the vessels’ corresponding order. The code used to generate the morphometric trees and solve for the hemodynamics is provided as Extended Data Data File S1.

As there is limited data on the organization of murine trees, morphometric information was taken from the literature and key parameters were fit to capture observed experimental data in the animal model^48, 50^. For the connectivity matrix, morphometric data from rats were adjusted to reduce the total number of vessel elements by removing the last row and column, leaving *N* = 10 for the simulations herein, which left a more appropriate total vessel number in comparison to *ex vivo* imaging^48, 50^. The highest order vessel element was assumed to be the main pulmonary artery (MPA). Diameter measurements for the MPA of each subject were taken with B-mode ultrasound. Cardiac output (CO) from Doppler ultrasound was used to determine the input boundary condition on flow to the simulated tree. The outlet boundary conditions on pressure were prescribed using the value of LVEDP. Subject-specific diameter ratios *ϕ_d_* were then fit to capture the RVSP as determined from the pressure measurements on catheterization, which ensured the total vascular resistance of the generated morphometric tree matched the pulmonary vascular resistance (PVR) calculated from the experimental data as *PVR = (RVSP – LVEDP) / CO*. Length of the main PA was taken from literature values for mice and length ratio *ϕ_l_* was fixed to that of the rat, due to its lesser contribution to resistance and lack of contribution to wall shear stress calculation^48, 50^. Wall shear stress values from all vessels of each order were then averaged for analysis. The computational framework for WSS simulation in PA trees is provided with Extended Data Data File S1.

### Tissue Processing

Following the RVSP measurement, mice were sacrificed by cervical dislocation under anesthesia, and the heart and lungs were harvested after flushing the pulmonary circulation with cold PBS. The lungs were fixed with 10% v/v neutral buffered formalin (NBF), embedded in paraffin for sectioning, and routine histology and morphometric light microscopic and confocal imaging analyses carried out as previously described^46^. Right ventricular hypertrophy (RVH) was assessed as the blotted dry weight of the right ventricle (RV) relative to the weight of the left ventricle plus septum (LV+S).

### Statistical analysis

Values from multiple experiments are shown as arithmetical mean±SEM. A *P*-value of <0.05 was considered significant. The number of samples in each group, the statistical test used, and the statistical significance is indicated in the figures. Data were analyzed using Prism version 8.4 (Graphpad).

### Data Availability

The data that support this study are available from the corresponding author upon reasonable request. The code used to generate the morphometric trees and solve for the hemodynamics is available in the Extended Data. The pulmonary EC-targeted AAV2-ESGHGYF vector is available upon request under an MTA with the University Medical Center Hamburg-Eppendorf. CUT&RUN data files are deposited in the Gene Expression Omnibus (GEO) under accession number GSE250550 and will be made publicly available upon publication.

