## Extended Data for "High Shear Stress Reduces ERG Causing Endothelial-Mesenchymal Transition and Pulmonary Arterial Hypertension"

**Supplementary Tables S1, S2, S4, S5**

*(Note: Supplementary Table S3 is provided as a separate Excel fine)*

**Supplementary Figures S1 – S4**

***Extended Data File S1***

*Computational framework for WSS* simulation in PA

### Supplementary Tables

#### Supplementary Table S1: Demographic characteristics of donor controls

#### A. Donor controls whose lungs were used in the experiments with PAEC

| ID | Age (yr)-Gender) | Race | Ethnicity | Cause of Death |
|---|---|---|---|---|
| Donor 1 | 1-M | White | Non-Hispanic | Anoxia/Drowning |
| Donor 2 | 12-M | Unknown | Non-Hispanic | Motor vehicle accident - head trauma following rollover motor vehicle ejection |
| Donor 3 | 16-M | White | Non-Hispanic | Gunshot wound resulting in subarachnoid hemorrhage |
| Donor 4 | 25-M | White | Non-Hispanic | Intracranial hemorrhage |

#### B. Donor controls whose lungs were used in the experiment with lung sections

| ID | Age (yr)-Gender | Race | Ethnicity | Cause of Death |
|---|---|---|---|---|
| Donor 6 | 28-F | White | Non-Hispanic | Motor vehicle accident - anoxia |
| Donor 7 | 43-F | White | Unknown | Cerebrovascular/Stroke |
| Donor 8 | 30-M | White | Non-Hispanic | Motor vehicle accident - head trauma blunt injury |
| Donor 9 | 34-F | Asian | Non-Hispanic | Cerebrovascular/Stroke ICH |

**Supplementary Table S2: Clinical and demographic characteristics of PAH patients**

| ID | Age(yr)-Gender | Race | Ethnicity | Diagnosis | PAP (s/d/m) | PVR (WU) | 6MWD (m) | PAH Medications |
|---|---|---|---|---|---|---|---|---|
| CHD-1 | 16-F | Unknown | Hispanic or Latino | APAH-CHD VSD | 88/50/64 | 17.86 | 269 | Sld, Amb, Bs, iv Trp, iv Ep, Tada, inh Trp |
| CHD-2 | 32-F | White | Hispanic or Latino | APAH-CHD VSD | 104/44/68 | 9.82 | 356.7 | Sld, Tada, Amb, SC Trp, inh Trp, Bs, inh ilo |
| CHD-3 | 44-F | Black/AA | Non-Hispanic | APAH-CHD VSD | 118/15/39 | N/A | 472.4 | iv Trp, Tada, Mac, Bs, inh Trp |
| CHD-4 | 40-F | White | Non-Hispanic | APAH-CHD VSD | 150/65/93 | 23.14 | 280.4 | Bs, iv Ep |
| FPAH-1 | 33-F | Black/AA | Non-Hispanic | FPAH BMPR2 mutation | 75/33/48 | 15.57 | 326.1 | iv Ep, Bs, Sld, inh Trp |
| FPAH-2 | 33-F | White | Non-Hispanic | FPAH BMPR2 mutation) | 87/29/48 | 9.74 | 288 | Bs, iv Trp, Sld, iv Ep |
| IPAH-1 | 41-F | White | Non-Hispanic | IPAH | 75/43/55 | 9.84 | 472.4 | Sld, Bs, iv Ep |
| IPAH-2 | 25-M | White | Hispanic or Latino | IPAH | 65/15/36 | N/A | 510.5 | iv Ep, Sld, iv Trp |

**Legend:** Clinical and demographic data for PAH patients were obtained from catheterization study performed closest to transplantation. PAH Diagnosis: Pulmonary arterial hypertension (PAH) classification: APAH-CHD, PAH associated with congenital heart defect. VSD, ventricular septal defect. (F)PAH, familial PAH; (I)PAH, idiopathic PAH; PAP (s/d/m), pulmonary arterial pressure (systolic/diastolic/mean). PVR, Pulmonary vascular resistance. Data were obtained from catheterization study performed closest to transplantation. 6MWD, distance walked in six minutes. Data were obtained from study performed closest to transplantation. N/A, not available. PAH medications are listed according to total drug exposure during treatment period up to transplantation, not necessarily in combination. Drugs abbreviations: Am (Ambrisentan), Bs (Bosentan), Ep (Epoprostenol), ilo (iloprost), Mac (macitentan), Sld (Sildenafil), Tada (tadalafil), Trp (Treprostinil). iv (intravenous), inh, inhaled. AA, African American.

**Supplementary Table S3: Significantly enriched motifs in H3K27ac peaks increased under LSS vs HSS conditions.**

Table S3 is provided as an Excel file.

Legend: Significantly enriched motifs were identified by Homer in H3K27ac peaks increased under LSS vs HSS conditions. Motifs are ranked in order of statistical significance. The DNA-binding domain of the transcription factor for each motif is given under the Motif Group column.

### Supplementary Table S4: Antibodies

#### A. Antibodies used for Immunohistochemistry and Immunofluorescence

| Antibody | Dilution | Company | Identifier | Source | Sample |
|---|---|---|---|---|---|
| CDH5 | 1/100 | Santa Cruz | sc9989 | mouse | human PAEC |
| CDH5 | 1/400 | Abcam | ab33168 | rabbit | human PAEC |
| ACTA2 | 1/200 | sigma | A2547 | mouse | human PAEC |
| PECAM1 | 1/100 | Abcam | ab9498 | mouse | human PAEC |
| FSP1 (S100A4) | 1/200 | Abcam | ab124805 | rabbit | human PAEC |
| ERG | 1/100 | Santa Cruz | sc376293 | mouse | human PAEC |
| ERG | 1/100 | Santa Cruz | sc376293 | mouse | human PAEC, mouse and human lungs |
| vWF | 1/250 | Abcam | ab6994 | rabbit | human lungs |
| ACTA2 | 1/100 | Abcam | ab184675 | mouse | mouse lungs |
| RFP (tdTomato) | 1/500 | ROCKLAND | 600-401-379 | Rabbit | mouse lungs |
| Luciferase | 1/100 | Abcam | ab21176 | Rabbit | mouse lungs |
| MECA32 | 1/20 | DSHB | MECA-32 | Rat | mouse lungs |

#### B. Antibodies used for Western Immunoblotting

| Antibody | Dilution | Company | Identifier | Source | Sample |
|---|---|---|---|---|---|
| NOS3 | 1/500 | BD biosciences | 610296 | Mouse | human PAEC, mouse lungs |
| ACTB | 1/1000 | Santa Cruz | sc47778 | Mouse | human PAEC |
| PECAM1 | 1/400 | Abcam | ab9498 | Mouse | human PAEC |
| CDH5 | 1/400 | Santa Cruz | sc9989 | Mouse | human PAEC |
| CDH5 | 1/10000 | Abcam | ab33168 | Rabbit | mouse lungs |
| ACTA2 | 1/1000 | Sigma-Aldrich | A2547 | Mouse | human PAEC |

| Antibody | Dilution | Company | Identifier | Source | Sample |
|---|---|---|---|---|---|
| SNAI1+SNAI2 | 1/1000 | Abcam | ab180714 | Rabbit | human PAEC |
| BMPR2 | 1/200 | BD biosciences | 612292 | Mouse | human PAEC, mouse lungs |
| KLF4 | 1/200 | R&D systems | AF3640 | Goat | human PAEC |
| ERG | 1/1000 | Abcam | ab92513 | Rabbit | human PAEC, mouse lungs |
| GAPDH | 1/1000 | Santa Cruz | sc25778 | Rabbit | mouse lungs |

### Supplementary Table S5: RT-qPCR Primers

| Human | Forward primer | Reverse primer |
|---|---|---|
| *NOS3* | TGATGGCGAAGCGAGTGAAG | ACTCATCCATACACAGGACCC |
| *B2M* | TTCTGGCCTGGAGGCTATC | TCAGGAAATTTGACTTTCCATTC |
| *PECAM1* | GCAACACAGTCCAGATAGTCGT | GACCTCAAACTGGGCATCAT |
| *CDH5* | GTTCACCTTCTGCGAGGATA | GTAGCTGGTGGTGTCCATCT |
| *ACTA2* | CCCTGAAGTACCCGATAGAACA | GGCAACACGAAGCTCATTG |
| *FSP1 (S100A4)* | GATGAGCAACTTGGACAGCAA | CTGGGCTGCTTATCTGGGAAG |
| *SNAI1* | CGAGTGGTTCTTCTGCGCTA | CTGCTGGAAGGTAAACTCTGGA |
| *SNAI2* | TGGTTGCTTCAAGGACACAT | GTTGCAGTGAGGGCAAGAA |
| *BMPR2* | CTGCGGCTGCTTCGCAGAAT | TGGTGTTGTGTCAGGAGGTGG |
| *KLF2* | CATCTGAAGGCGCATCTG | CGTGTGCTTTCGGTAGTGG |
| *KLF4* | GGGAGAAGACACTGCGTCA | GGAAGCACTGGGGGAAGT |
| *ERG* | GCCAGGTGAATGGCTCAA | AGTTCATCCCAACGGTGTC |
| *ACTB* | CCAACCGCGAGAAGATGA | CCAGAGGCGTACAGGGATAG |

### Supplementary Figures and Legends:

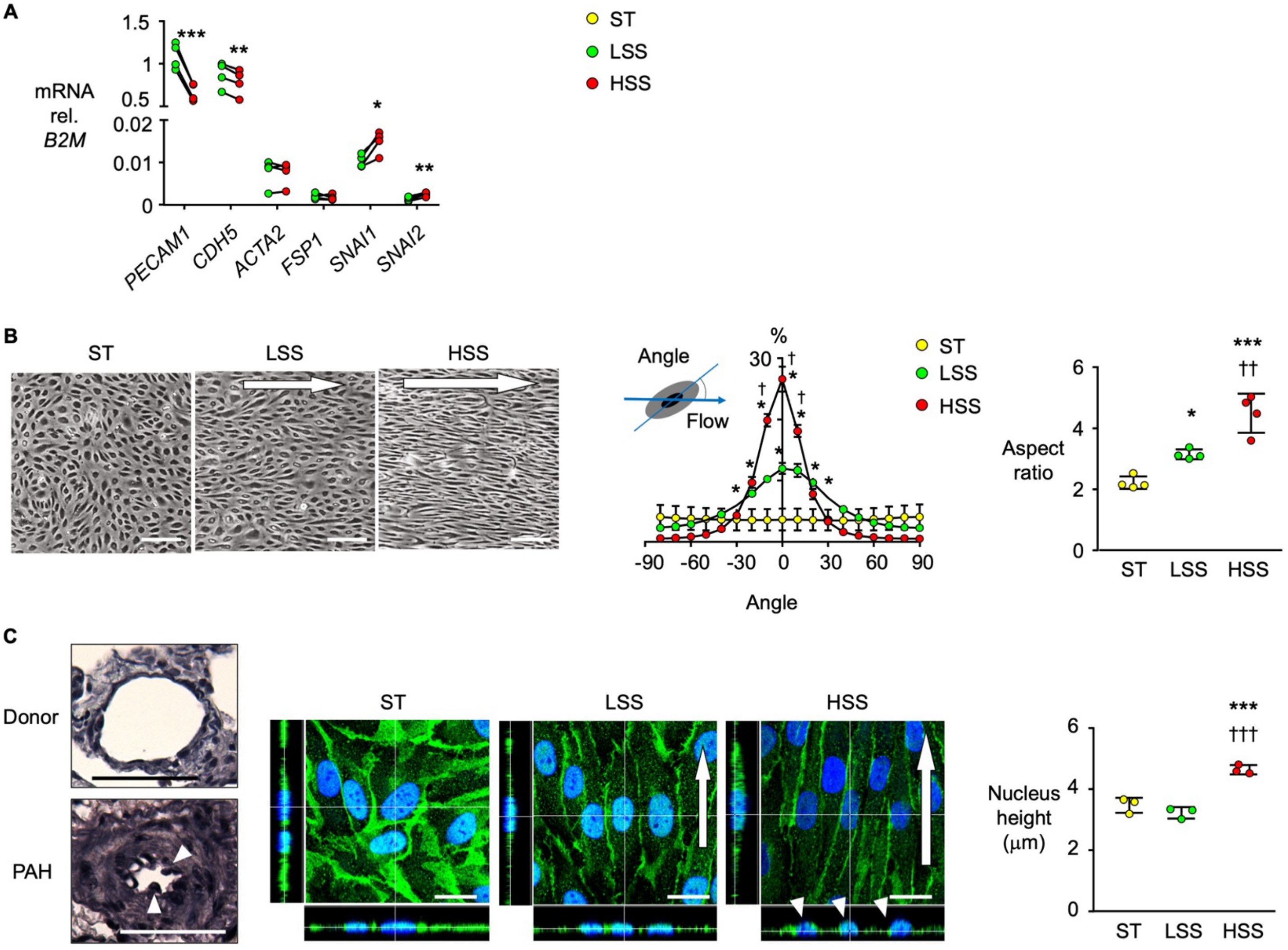


**Supplementary Figure S1: HSS increases alignment and induces morphologic changes.** Human donor PAEC were exposed to 24 h of high shear stress (HSS, 100 dyn/cm²) or 24 h of laminar shear stress (LSS, 15 dyn/cm²) after preconditioning by 24 h of LSS and compared to cells cultured for 48 h without flow (static condition, ST). **(A)** mRNA levels of EC markers (PECAM1, CDH5), Mesenchymal cell markers (ACTA2, FSP1), and EndMT inducing transcription factors (SNAI1, SNAI2) in PAEC under LSS and HSS, assessed by RT-qPCR. Individual data points are shown, with n=4 biological replicates. *p<0.05, **p<0.01, ***p<0.001 vs. LSS, by paired Student t test. (**B**) Left, Representative phase-contrast light microscopic images of PAEC under static, LSS and HSS. Scale bar: 100 µm. Arrow indicates direction of flow. Middle, quantification of cell orientation relative to flow direction, and percentage of frequency, with quantification. Right, spindle shape morphology, calculated by the ratio of the longer axis length relative to the shorter axis length. n=4 biological replicates. **(C)** Left, representative elastic van Gieson staining of distal PAs. Top: Flattened-nucleus of PAEC of Control Donor. Bottom: protruding-nucleus of PAEC of VSD patients (arrowhead), Scale bar: 50 µm. Middle: representative confocal 3D image of PAEC stained for CDH5 (green), and DAPI (blue). HSS induces the protruding-nucleus (arrowhead) compared to LSS and ST. Scale bar, 20 µm. Arrow indicates direction of flow. Right, quantification of nucleus height. n=3 biological replicates. In (**B, C**), data shown as Mean±SEM. *p<0.05, ***p<0.001 vs. Static, $^{\dagger}$p<0.05, $^{\dagger\dagger}$p<0.01, $^{\dagger\dagger\dagger}$p<0.001 vs. LSS by one-way ANOVA followed by Tukey's multiple comparisons test.

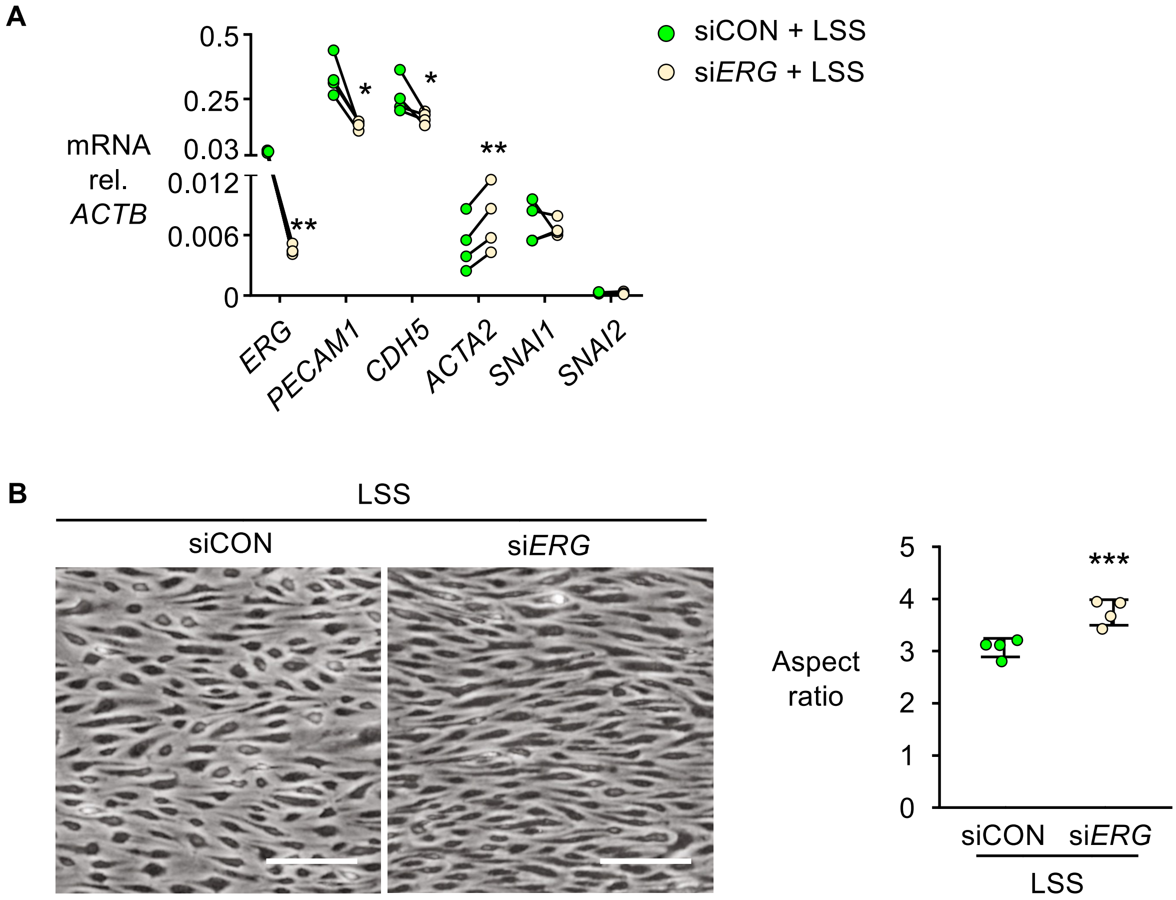


**Supplementary Figure S2: Reduced *ERG* under LSS leads to spindle-shape morphology and gene expression changes consistent with EndMT.**

PAEC were transfected with siRNA targeting *ERG* (si*ERG*) or with nontargeting siRNA (siCON) and cultured under laminar shear stress (LSS, 15 dyn/cm$^2$) for 48 hr. **(A)** mRNA levels of *ERG, PECAM1, CDH5, ACTA2, SNAI1* and *SNAI2*, assessed by RT-qPCR. Individual data points are shown for n=4 biological replicates. *p<0.05; **p<0.01 vs. siCON under LSS, by paired Student t-test. **(B)** Representative phase-contrast light microscopic images of the PAEC, with quantification of the spindle shape morphology on the right. Data shown as mean±SEM of n=4 biological replicates. ***p<0.001 vs. siCON by Student t-test.

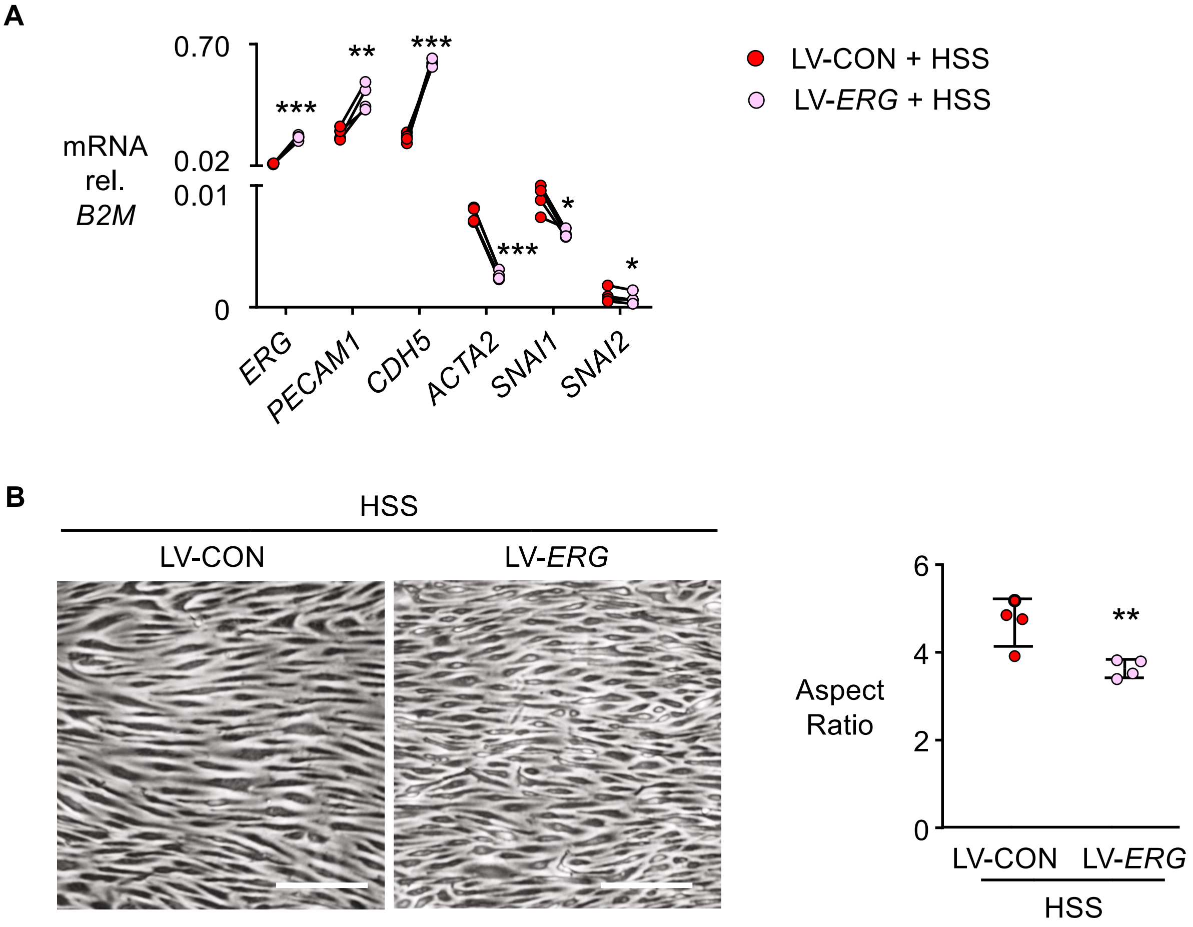


**Supplementary Figure S3: Replenishing *Erg* in PAEC under HSS induces changes in gene expression consistent with reversal of EndMT and rescues the spindle shape.**

PAEC were transfected with by lentiviral vector encoding *ERG* (LV-*ERG*) or with control LV encoding *EGFP* (LV-CON) vector, and subjected to high shear stress (HSS, 100 dyn/cm$^2$) 24 h, after preconditioning by 24 h of laminar shear stress (15 dyn/cm$^2$). **(A)** Gene expression levels of *ERG, PECAM1, CDH5, ACTA2*, *SNAI1* and *SNAI2*, assessed by RT-qPCR. **(B)** Representative phase-contrast light microscopic images, with quantification of the spindle shape morphology on the right. Scale bar: 100 µm.

Individual data points are shown, with mean±SEM of n=4 biological replicates. *p<0.05, **p<0.01, ***p<0.001 vs. LV-CON by paired Student t test.

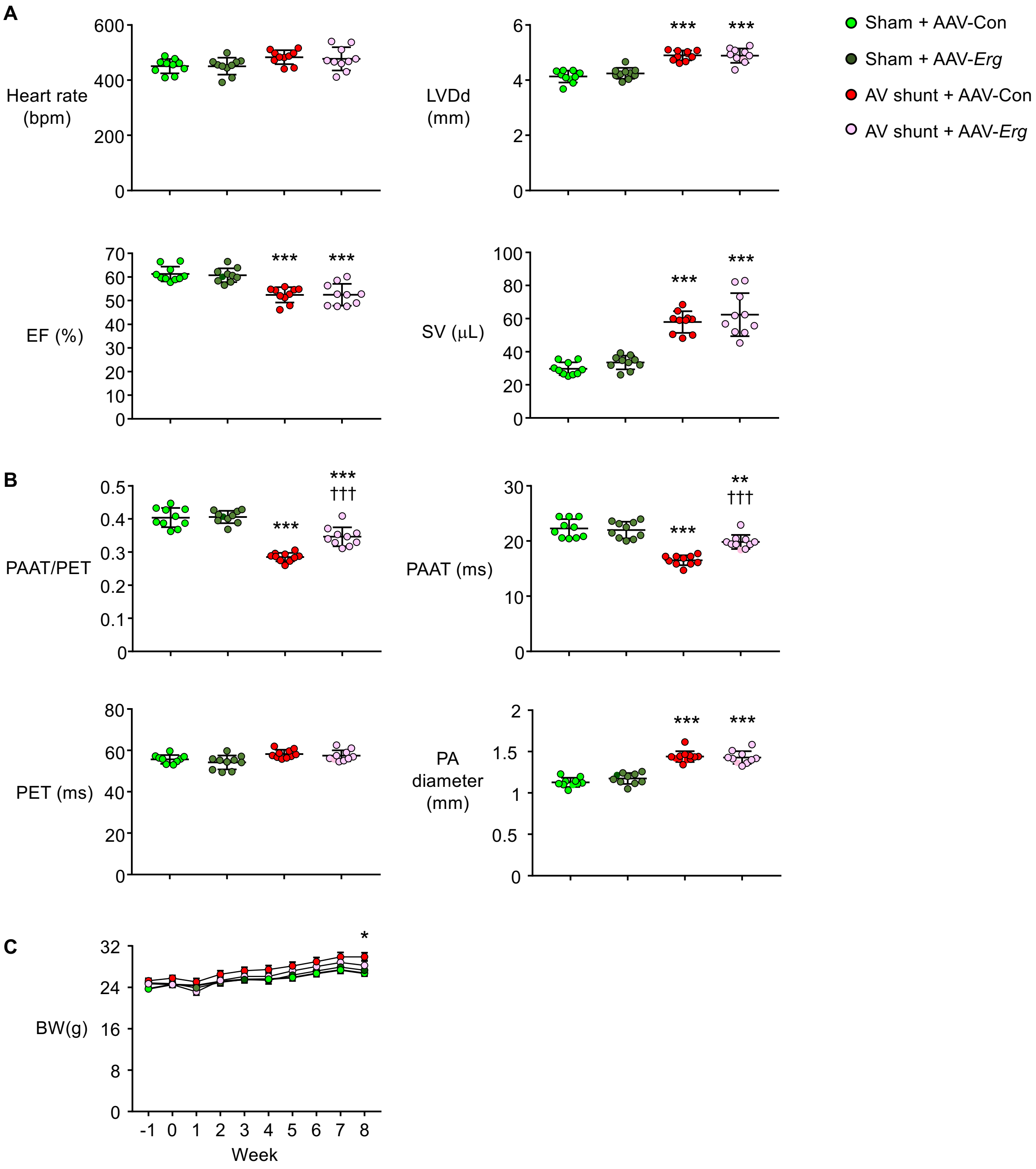


**Supplementary Figure S4: Echocardiographic Features**

Mice underwent AV shunt or sham operation and were treated with AAV vectors encoding *Erg* (AAV-*Erg)* or luciferase (AAV-Con) under the experimental protocol described in Figure 5. Hemodynamic parameters were assessed at the 8-week timepoint, as described in "Methods". **(A)** Heart rate, left ventricular (LV) diameter at the end diastoles (LVDd), ejection fraction (EF) and stroke volume (SV). **(B)** Pulmonary acceleration time per pulmonary ejection time

(PAAT/PET), pulmonary acceleration time (PAAT), pulmonary ejection time (PET), and PA diameter. **(C)** Body weight (BW) of the four experimental groups, measured weekly from week -1 to the end of the $8^{th}$ week.

In **(A, B),** each point represents a mouse, and the mean±SEM shown. In **(C)**, each point represents the mean±SEM of n=10 mice per group. *p<0.05, **p<0.01, ***p<0.001 vs. respective Sham group, and $^{†††}$p<0.001 vs. AV shunt with luciferase group (AAV-Con), by one-way ANOVA followed by Tukey's multiple comparisons test.
